# Novel tetraplex qPCR assays for simultaneous detection and identification of *Xylella fastidiosa* subspecies in plant tissues

**DOI:** 10.1101/699371

**Authors:** Dupas Enora, Briand Martial, Jacques Marie-Agnès, Cesbron Sophie

**Author notes:** **Correspondance:** Sophie Cesbron, Marie-Agnès Jacques.

## Abstract

*Xylella fastidiosa* is an insect-borne bacterium confined to the xylem vessels of plants. This plant pathogen has a broad host range estimated to 560 plant species. Five subspecies of the pathogen with different but overlapping host ranges have been described, but only three subspecies are widely accepted, namely subspecies *fastidiosa*, *multiplex* and *pauca*. Initially limited to the Americas, *Xf* has been detected in Europe since 2013. As management of *X. fastidiosa* outbreaks in Europe depends on the identification of the subspecies, accurate determination of the subspecies in infected plants as early as possible is of major interest. Thus, we developed various tetraplex and triplex qPCR assays for *Xylella fastidiosa* detection and subspecies identification *in planta* in a single reaction. We designed primers and probes using SkIf, a bioinformatics tool based on *k*-mers, to detect specific signatures of the species and subspecies from a dataset of 58 genome sequences representative of *X. fastidiosa* diversity. We tested the qPCR assays on 39 target and 30 non-target strains, as well as on 13 different plant species spiked with strains of the different subspecies of *X. fastidiosa*, and on samples from various environmental and inoculated host plants. Sensitivity of simplex assays was equal or slightly better than the reference protocol on purified DNA. Tetraplex qPCR assays had the same sensitivity than the reference protocol and allowed *X. fastidiosa* detection in all spiked matrices up to 10^3^ cells.mL^−1^. Moreover, mix infections of two to three subspecies could be detected in the same sample with tetraplex assays. In environmental plant samples, the tetraplex qPCR assays allowed subspecies identification when the current method based on multilocus sequence typing failed. The qPCR assays described here are robust and modular tools that are efficient for differentiating *X. fastidiosa* subspecies directly in plant samples.

## 1 Introduction

*Xylella fastidiosa* (*Xf*) is a worldwide insect-transmitted plant pathogenic bacterium that presents a very large host range. Altogether, 563 plant species grouped into 82 botanical families have been reported as *Xf* hosts (EFSA, 2018a). Plants with a major socio-economic interest such as grapevine, citrus, coffee, and olive trees are hosts of *Xf* (EFSA, 2018a). Forest trees, shade trees, ornamentals and landscape species are included in the host plant database making this pathogen a potential worldwide threat (EFSA, 2018a). Disease management of *Xf* is impeded by its asymptomatic period that can last several years (EFSA, 2018b).

This bacterial species is genetically diverse as five subspecies including *fastidiosa, morus, multiplex, pauca* and *sandyi* are currently described (EFSA, 2018b). Although this subspecies delineation was initially associated to *Xf* host range and places of occurrence, more and more observations report infection of a given host by various subspecies (Denancé *et al*., 2017; EPPO, 2018b; Denancé et al., 2019; Nunney *et al*., 2019). Homologous recombination events were detected in Xf and were suspected to be associated with host-shift, as documented for the subspecies morus (Nunney et al., 2014). But intrasubspecific homologous recombination events could be more frequent than intersubspecific events (Potnis et al., 2019). Based on genome sequence analyses, it was proposed to merge the subspecies *fastidiosa, morus* and *sandyi* in the subspecies *fastidiosa* (hereafter referred to *Xff sensu lato* (*Xffsl*) to avoid confusion with classical *Xff*), the subspecies *multiplex* and *pauca* remaining coherent groups and distantly related from *Xff* (Marcelletti and Scortichini, 2016; Denancé *et al*., 2019). The method generally used to identify strains at the subspecies level is based on the sequencing of seven housekeeping genes (*cysG, gltT, holC, leuA, malF, nuoL* and *petC)* of the dedicated MultiLocus Sequence Typing (MLST) scheme (Yuan *et al*., 2010).

In Europe, *Xf* has been reported for the first time in Apulia area, Italy, in olive trees (Saponari *et al*., 2013). Then, *Xf* was detected in 2015 in France, more precisely in Corsica and in the French Riviera region, mainly on *Polygala myrtifolia* and other ornamentals (Denancé *et al*., 2017). Two years later, *Xf* has been reported in the Balearic Islands mostly in olive tree, grapevine and sweet cherry and in continental Spain in almond trees (Landa, 2017). More recently, in October 2018, the presence of *X. fastidiosa* subsp. *multiplex* was reported in Monte Argentario (Tuscany, Italy), and in January 2019 the subsp. *multiplex* was identified in Portugal (region of Porto), and both reports concerned ornamentals (EPPO, 2019). Since the first report, four subspecies, *fastidiosa, multiplex, pauca* and *sandyi* have been identified in Europe (Jacques et al., 2016; Denancé et al., 2017; Cruaud et al., 2018). A number of cases of imported plants being infected by *Xf* has also been reported in Europe since 2012 (EPPO, 2019). Being present in Europe, *Xf* was first listed as an A1 regulated pathogen. *Xf* is now reported in the Annex I/A2 of the directive 2000/29/CE and in the EPPO A2 list (C/2017/4883, 2017; EPPO, 2018a).

Apart the sympatry of several subspecies at the local, regional or state level, cases of mix infection of plants have been described. In 2005 in California, an almond tree has been reported infected by two types of *Xf* strains, revealing the first case of mix infection by *Xf* (Chen *et al*., 2005). Recently, in coffee trees imported into Europe from Central America, the MLST revealed a mix infection with two different sequence types (STs) of *Xf* from two subspecies: *pauca* and *fastidiosa* (Bergsma-Vlami *et al*., 2017). In France, a *Polygala myrtifolia* plant was found mix infected with strains of two different STs (Denancé *et al*., 2017). Reported cases of undetermined sequences of housekeeping gene alleles was an indication of mix infections in plants (Denancé *et al*., 2017).

Because in Europe the subspecies identification is necessary to set up outbreak management, it is of major interest to have access to reliable tools for the detection and identification of *Xf*. As *Xf* isolation is tedious, detection and identification of subspecies are performed directly on plant extracts (Denancé *et al*., 2017). To date, tests based on loop-mediated isothermal amplification (LAMP) (Harper *et al*., 2010), conventional PCR (Minsavage *et al*., 1994; Hernandez-Martinez *et al*., 2006), and quantitative PCR (qPCR) (Francis *et al*., 2006; Harper *et al*., 2010; Li *et al*., 2013; Ouyang *et al*., 2013) targeting specific regions at the species or subspecies level are available. Among these tests, the qPCR assay developed by Harper *et al.* (2010) has been identified as one of the most appropriate for the detection of *Xf*, as it has shown a high diagnostic sensitivity compared to others qPCR assays, detects all subspecies, has no cross-reactivity with any other bacterial species and has been successfully used on a wide range of plants (Modesti *et al*., 2017; Reisenzein, 2017). Several tests have been proposed to identify one or more subspecies but no test is currently available to identify all subspecies. The subspecies identification is then routinely performed by MLST, but this method while accurate and portable is time consuming, labor intensive and expensive. From 2018, sequences of only two housekeeping genes (*rpoD* and *cysG* or *rpoD* and *malF*) are required for subspecies identification in France, while other sets of gene pairs are recommended by EPPO (EPPO, 2018b).

In recent years, multiplexed Taqman qPCR has become a useful tool for the identification and quantification of pathogens in different areas such as food safety (Köppel *et al*., 2019; Wei *et al*., 2019), medical environment (Janse *et al*., 2013; Kamau *et al*., 2013), agronomics (Wei *et al*., 2008; Zitnick-Anderson *et al*., 2018), GMO detection (Choi *et al*., 2018; Wang *et al*., 2018), and the environment (Hulley *et al*., 2019). For plant pathogens these methods have been tested on samples of naturally infected plants, spiked samples (Li *et al*., 2009; Willsey *et al*., 2018), and on mixtures of plant and pathogen DNAs (Abraham *et al*., 2018). *Xf*-specific multiplexed qPCR assays have already been developed based on the combination of primers designed by Harper *et al*. (2010) and Ouyang *et al*. (2013) (Bonants *et al*., 2018). Other tests were proposed to differentiate *Xf* from phytoplasmas sharing common host plants (Ito and Suzaki, 2017) and to differentiate the subspecies *fastidiosa* from the subspecies *multiplex* (Burbank and Ortega, 2018). However, none of them allows the differential identification of all *Xf* subspecies.

In this study, we described the development and evaluation of six multiplex qPCR assays for the detection and identification of *Xf* subspecies. These tests have been designed and tested *in silico* on a wide range of target and non-target genomic sequences, *in vitro* on target and non-target bacterial strains, on *Xf*-spiked plant extracts, and finally *in planta* on samples from environmental or inoculated plants. These assays allowed the detection of *Xf* subspecies up to 10 pg.mL^−1^ of DNA, 1×10^3^ CFU.mL^−1^ in spiked samples and allow the identification of *Xf* subspecies in environmental plant samples that cannot be typed using MLST. These multiplex qPCR assays offer a new, faster, more reliable, more specific, more sensitive, and less expensive tool than MLST.

## 2 Materials and methods

### 2.1 Bacterial strains and growth conditions

Collections of 39 strains representing the different *Xf* subspecies, 28 strains from other plant-pathogenic bacterium genera (*Agrobacterium*, *Clavibacter*, *Dickeya*, *Erwinia*, *Pantoea*, *Pseudomonas*, *Stenotrophomonas*, *Xanthomonas* and *Xylophilus*), and two strains from plant endosymbionts (*Ensifer* and *Rhizobium*) were used (Table 1). A set of 12 *Xf* strains of the subsp. *multiplex* and one strain of the subsp. *sandyi* were kindly provided by Leonardo De la Fuente (Auburn University, AL, USA). The other 57 strains were provided by the French Collection of Plant-Associated Bacteria (CIRM-CFBP; https://www6.inra.fr/cirm_eng/CFBP-Plant-Associated-Bacteria). *Xf* strains were grown on BCYE (Wells *et al*., 1981) or modified PWG media (agar 12 g.L^−1^; soytone 4 g.L^−1^; bacto tryptone 1 g.L^−1^; MgSO_4_.7H_2_O 0,4 g.L^−1^; K_2_HPO_4_ 1.2 g.L^−1^; KH_2_PO_4_ 1 g.L^−1^; hemin chloride (0.1% in NaOH 0.05 M) 10 ml.L^−1^; BSA (7.5%) 24 ml.L^−1^; L-glutamine 4 g.L^−1^) at 28°C for one to two weeks. Other strains were grown at 25°C for one to two days on: MG media (Mougel *et al*., 2001) for *Agrobacterium* and *Rhizobium*, TSA (tryptone soy broth 30 g.L^−1^; agar 15 g.L^−1^) for *Clavibacter*, *Ensifer*, *Stenotrophomonas*, *Xanthomonas* and *Xylophilus* and King’s B medium (KH_2_PO_4_ 1.5 g.L^−1^; MgSO_4_, 7H_2_O 1.5 g.L^−1^; protease peptone 20 g.L^−1^, glycerol 10 mL.L-1; agar 15 g.L^−1^) for *Dickeya*, *Erwinia*, *Pantoea* and *Pseudomonas*. For qPCR assays, bacterial suspensions were prepared from fresh cultures in sterile distilled water, adjusted at OD_600nm_ = 0.1. To evaluate assay specificity bacterial suspensions were boiled for 20 min, followed by a thermal shock on ice and a centrifugation at 10,000 g during 10 min.

**Table 1:**
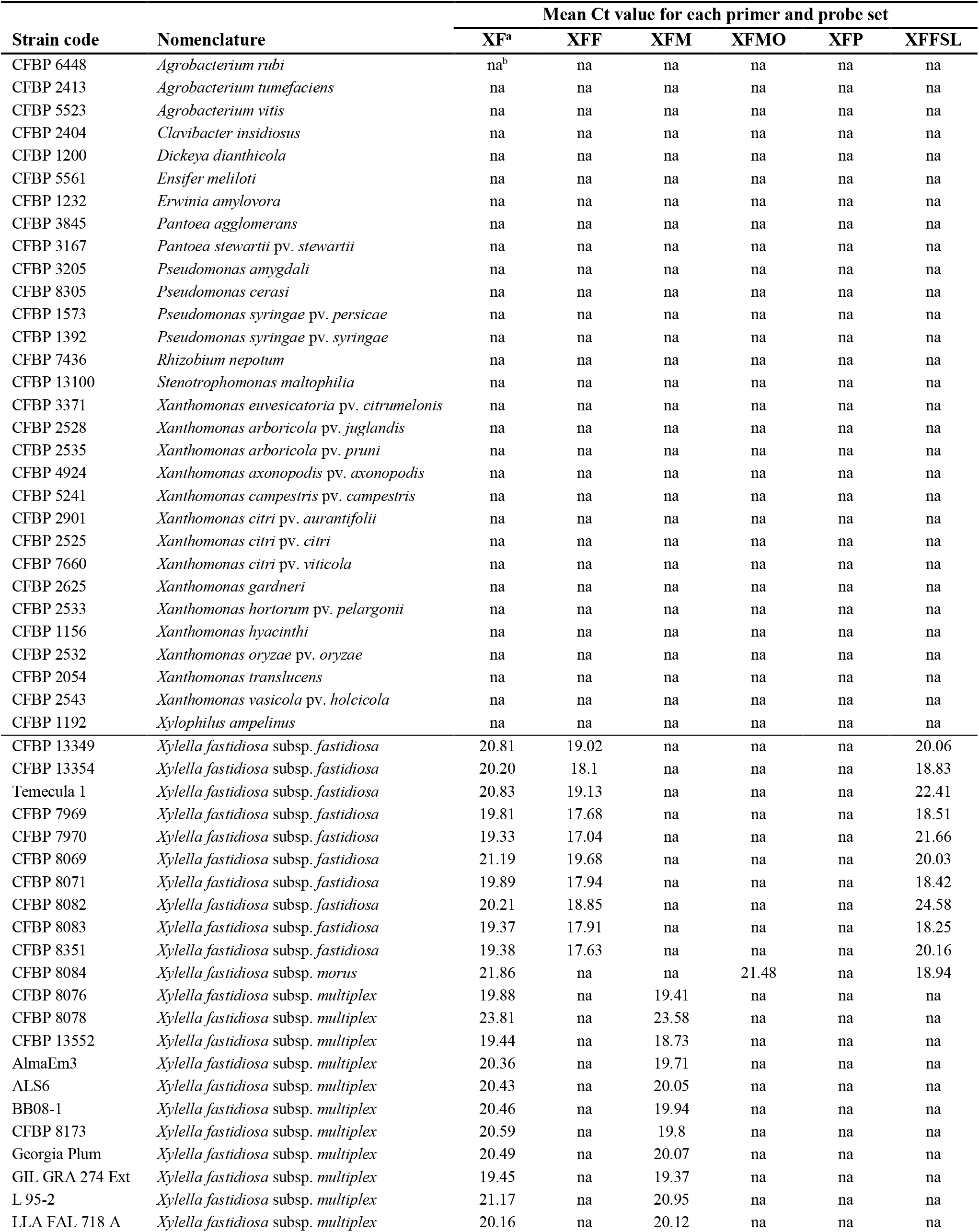
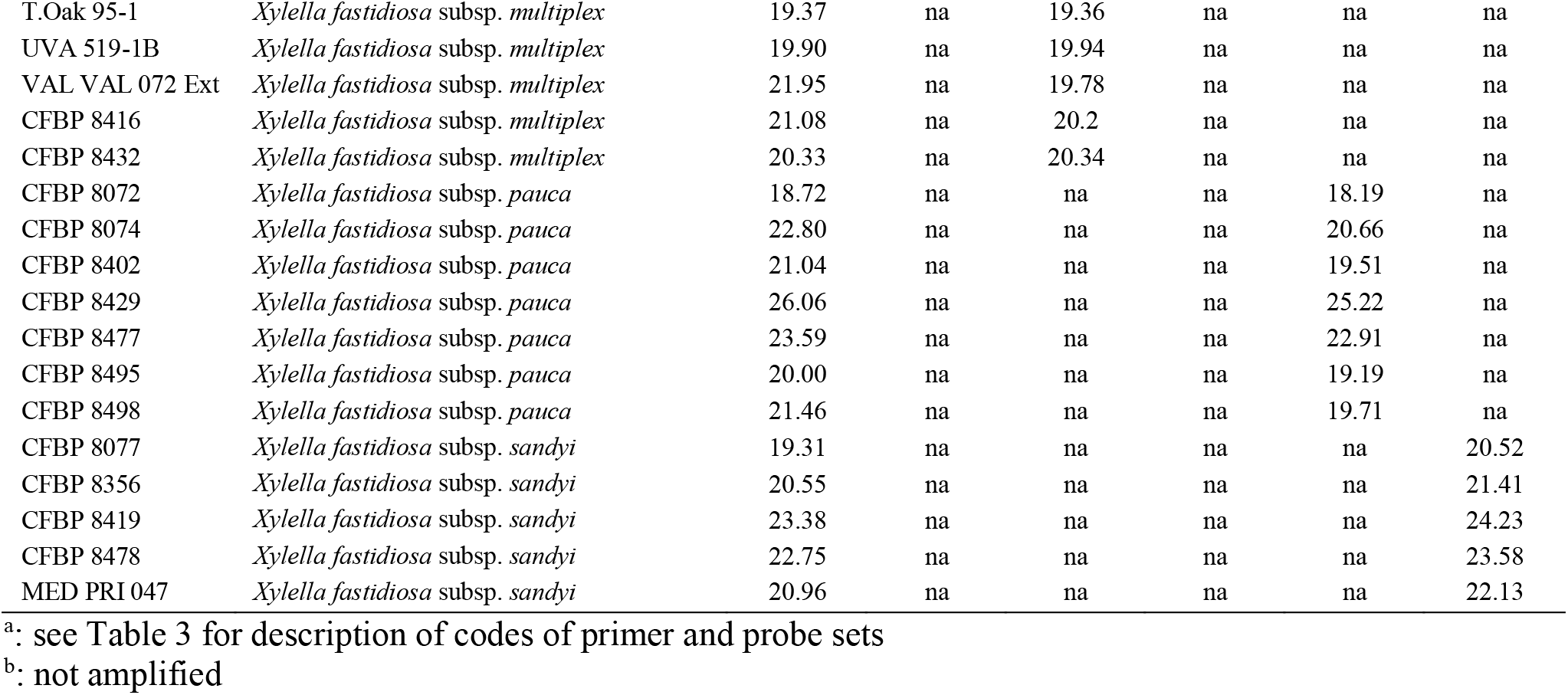
List of strains used in this study and signals obtained with the primers and probe combinations in simplex qPCR assays on DNA suspensions calibrated at OD_600nm_ = 0.1.

| Strain code | Nomenclature | Mean Ct value for each primer and probe set |  |  |  |  |  |
| --- | --- | --- | --- | --- | --- | --- | --- |
|  |  | XF<sup>a</sup> | XFF | XFM | XFMO | XFP | XFFSL |
| CFBP 6448 | *Agrobacterium rubi* | na<sup>b</sup> | na | na | na | na | na |
| CFBP 2413 | *Agrobacterium tumefaciens* | na | na | na | na | na | na |
| CFBP 5523 | *Agrobacterium vitis* | na | na | na | na | na | na |
| CFBP 2404 | *Clavibacter insidiosus* | na | na | na | na | na | na |
| CFBP 1200 | *Dickeya dianthicola* | na | na | na | na | na | na |
| CFBP 5561 | *Ensifer meliloti* | na | na | na | na | na | na |
| CFBP 1232 | *Erwinia amylovora* | na | na | na | na | na | na |
| CFBP 3845 | *Pantoea agglomerans* | na | na | na | na | na | na |
| CFBP 3167 | *Pantoea stewartii* pv. *stewartii* | na | na | na | na | na | na |
| CFBP 3205 | *Pseudomonas amygdali* | na | na | na | na | na | na |
| CFBP 8305 | *Pseudomonas cerasi* | na | na | na | na | na | na |
| CFBP 1573 | *Pseudomonas syringae* pv. *persicae* | na | na | na | na | na | na |
| CFBP 1392 | *Pseudomonas syringae* pv. *syringae* | na | na | na | na | na | na |
| CFBP 7436 | *Rhizobium nepotum* | na | na | na | na | na | na |
| CFBP 13100 | *Stenotrophomonas maltophilia* | na | na | na | na | na | na |
| CFBP 3371 | *Xanthomonas euvesicatoria* pv. *citrumelonis* | na | na | na | na | na | na |
| CFBP 2528 | *Xanthomonas arboricola* pv. *juglandis* | na | na | na | na | na | na |
| CFBP 2535 | *Xanthomonas arboricola* pv. *pruni* | na | na | na | na | na | na |
| CFBP 4924 | *Xanthomonas axonopodis* pv. *axonopodis* | na | na | na | na | na | na |
| CFBP 5241 | *Xanthomonas campestris* pv. *campestris* | na | na | na | na | na | na |
| CFBP 2901 | *Xanthomonas citri* pv. *aurantifolii* | na | na | na | na | na | na |
| CFBP 2525 | *Xanthomonas citri* pv. *citri* | na | na | na | na | na | na |
| CFBP 7660 | *Xanthomonas citri* pv. *viticola* | na | na | na | na | na | na |
| CFBP 2625 | *Xanthomonas gardneri* | na | na | na | na | na | na |
| CFBP 2533 | *Xanthomonas hortorum* pv. *pelargonii* | na | na | na | na | na | na |
| CFBP 1156 | *Xanthomonas hyacinthi* | na | na | na | na | na | na |
| CFBP 2532 | *Xanthomonas oryzae* pv. *oryzae* | na | na | na | na | na | na |
| CFBP 2054 | *Xanthomonas translucens* | na | na | na | na | na | na |
| CFBP 2543 | *Xanthomonas vasicola* pv. *holcicola* | na | na | na | na | na | na |
| CFBP 1192 | *Xylophilus ampelinus* | na | na | na | na | na | na |
| CFBP 13349 | *Xylella fastidiosa* subsp. *fastidiosa* | 20.81 | 19.02 | na | na | na | 20.06 |
| CFBP 13354 | *Xylella fastidiosa* subsp. *fastidiosa* | 20.20 | 18.1 | na | na | na | 18.83 |
| Temecula 1 | *Xylella fastidiosa* subsp. *fastidiosa* | 20.83 | 19.13 | na | na | na | 22.41 |
| CFBP 7969 | *Xylella fastidiosa* subsp. *fastidiosa* | 19.81 | 17.68 | na | na | na | 18.51 |
| CFBP 7970 | *Xylella fastidiosa* subsp. *fastidiosa* | 19.33 | 17.04 | na | na | na | 21.66 |
| CFBP 8069 | *Xylella fastidiosa* subsp. *fastidiosa* | 21.19 | 19.68 | na | na | na | 20.03 |
| CFBP 8071 | *Xylella fastidiosa* subsp. *fastidiosa* | 19.89 | 17.94 | na | na | na | 18.42 |
| CFBP 8082 | *Xylella fastidiosa* subsp. *fastidiosa* | 20.21 | 18.85 | na | na | na | 24.58 |
| CFBP 8083 | *Xylella fastidiosa* subsp. *fastidiosa* | 19.37 | 17.91 | na | na | na | 18.25 |
| CFBP 8351 | *Xylella fastidiosa* subsp. *fastidiosa* | 19.38 | 17.63 | na | na | na | 20.16 |
| CFBP 8084 | *Xylella fastidiosa* subsp. *morus* | 21.86 | na | na | 21.48 | na | 18.94 |
| CFBP 8076 | *Xylella fastidiosa* subsp. *multiplex* | 19.88 | na | 19.41 | na | na | na |
| CFBP 8078 | *Xylella fastidiosa* subsp. *multiplex* | 23.81 | na | 23.58 | na | na | na |
| CFBP 13552 | *Xylella fastidiosa* subsp. *multiplex* | 19.44 | na | 18.73 | na | na | na |
| AlmaEm3 | *Xylella fastidiosa* subsp. *multiplex* | 20.36 | na | 19.71 | na | na | na |
| ALS6 | *Xylella fastidiosa* subsp. *multiplex* | 20.43 | na | 20.05 | na | na | na |
| BB08-1 | *Xylella fastidiosa* subsp. *multiplex* | 20.46 | na | 19.94 | na | na | na |
| CFBP 8173 | *Xylella fastidiosa* subsp. *multiplex* | 20.59 | na | 19.8 | na | na | na |
| Georgia Plum | *Xylella fastidiosa* subsp. *multiplex* | 20.49 | na | 20.07 | na | na | na |
| GIL GRA 274 Ext | *Xylella fastidiosa* subsp. *multiplex* | 19.45 | na | 19.37 | na | na | na |
| L 95-2 | *Xylella fastidiosa* subsp. *multiplex* | 21.17 | na | 20.95 | na | na | na |
| LLA FAL 718 A | *Xylella fastidiosa* subsp. *multiplex* | 20.16 | na | 20.12 | na | na | na |
| T.Oak 95-1 | *Xylella fastidiosa* subsp. *multiplex* | 19.37 | na | 19.36 | na | na | na |
| UVA 519-1B | *Xylella fastidiosa* subsp. *multiplex* | 19.90 | na | 19.94 | na | na | na |
| VAL VAL 072 Ext | *Xylella fastidiosa* subsp. *multiplex* | 21.95 | na | 19.78 | na | na | na |
| CFBP 8416 | *Xylella fastidiosa* subsp. *multiplex* | 21.08 | na | 20.2 | na | na | na |
| CFBP 8432 | *Xylella fastidiosa* subsp. *multiplex* | 20.33 | na | 20.34 | na | na | na |
| CFBP 8072 | *Xylella fastidiosa* subsp. *pauca* | 18.72 | na | na | na | 18.19 | na |
| CFBP 8074 | *Xylella fastidiosa* subsp. *pauca* | 22.80 | na | na | na | 20.66 | na |
| CFBP 8402 | *Xylella fastidiosa* subsp. *pauca* | 21.04 | na | na | na | 19.51 | na |
| CFBP 8429 | *Xylella fastidiosa* subsp. *pauca* | 26.06 | na | na | na | 25.22 | na |
| CFBP 8477 | *Xylella fastidiosa* subsp. *pauca* | 23.59 | na | na | na | 22.91 | na |
| CFBP 8495 | *Xylella fastidiosa* subsp. *pauca* | 20.00 | na | na | na | 19.19 | na |
| CFBP 8498 | *Xylella fastidiosa* subsp. *pauca* | 21.46 | na | na | na | 19.71 | na |
| CFBP 8077 | *Xylella fastidiosa* subsp. *sandyi* | 19.31 | na | na | na | na | 20.52 |
| CFBP 8356 | *Xylella fastidiosa* subsp. *sandyi* | 20.55 | na | na | na | na | 21.41 |
| CFBP 8419 | *Xylella fastidiosa* subsp. *sandyi* | 23.38 | na | na | na | na | 24.23 |
| CFBP 8478 | *Xylella fastidiosa* subsp. *sandyi* | 22.75 | na | na | na | na | 23.58 |
| MED PRI 047 | *Xylella fastidiosa* subsp. *sandyi* | 20.96 | na | na | na | na | 22.13 |

<sup>a</sup>: see Table 3 for description of codes of primer and probe sets
<sup>b</sup>: not amplified

### 2.2 Plant material

Petioles or midribs were collected in 2018 from healthy plants of 13 species (*Helichrysum italicum, Lavandula angustifolia, Nerium oleander, Olea europaea, Prunus cerasus, Prunus dulcis, Quercus ilex, Quercus robur* and *Rosmarinus officinalis*) growing in orchards adjacent to INRA center or purchased in nurseries (*Vitis vinifera, Citrus clementina* and *Polygala myrtifolia*). These species are either not known to be host of *Xf* in France or were sampled in *Xf*-free areas. Symptomless *Cistus monspeliensis* plant material was collected in Corsica outside any recorded *Xf*-focus by the National Botanical Conservatory of Corsica (CNBC).

Plants were collected in June 2017 and in October 2018 in Corsica, France, based on symptoms and were pre-tested using a modified extraction procedure based on CTAB and/or QuickPickTM SML Plant DNA Kit (Bio-Nobile, Turku, Finland) as described in PM7/24 (EPPO, 2018b). Samples were first finely chopped and then sonicated (1 min, 42KHz) in a Branson apparatus. A 15 min incubation step at room temperature was performed before DNA extraction. The frozen DNA solutions of 20 greenhouse inoculated plant materials were used to evaluate the multiplex qPCR assays.

### 2.3 Production of inoculated plants

*X. fastidiosa* strains CFBP 7970 (*Xff*), CFBP 8077 (*Xfs*), CFBP 8402 (*Xfp*), CFBP 8416 (*Xfm*) and CFBP 8418 (*Xfm*) were inoculated in six month-old grafted plants of *Vitis vinifera* cv Chardonnay, *Vitis vinifera* cv Cabernet Franc, in 1.5 years-old grafted plants of *Prunus armeniaca* var Bergeron, *Olea europaea* cv Aglandau, *Olea europaea* cv Capanaccia, and *Olea europaea* cv Sabine. Plants were grown in a confined growth chamber at 24°C with 16 h of daylight and at 20°C during night, under 70% relative humidity. Plants were watered daily with water supplemented with 1.4 g.L^−1^ nitrogen:phosphorus:potassium fertilizer (16:8:32). Plants were inoculated by the needle puncture method. A 10 µL drop of inoculum calibrated at OD_600nm_ = 0.5 was placed on the node of a growing young stem and punctured with a needle. After six months for vines and apricot trees, and one year for olive trees, samples at the inoculation point were tested by the Harper’s qPCR test and typed using the classical *Xf* MLST scheme as described in Denancé et al. (2017). The samples were stored at −20°C before being analyzed. Plant inoculations were carried out under quarantine at IRHS, Centre INRA, Beaucouzé, France under the agreement no. 2013119-0002 from the Prefecture de la Région Pays de la Loire, France.

### 2.4 Spiking of samples and DNA extraction

Prior to DNA extraction, plant samples were inoculated by mixing 1 g of healthy plant material with 0.5 mL of a bacterial suspension, at a known concentration, and ground with 4.5 mL of sterile distilled water. Each matrix was spiked in order to end up with concentrations ranging from 1×10^6^ CFU.mL^−1^ to 10 CFU.mL^−1^. Spiking with more than one strain was done in equal amounts to end up with final concentrations ranging from 1×10^6^ CFU.mL^−1^ to 1×10 CFU.mL^−1^. Samples from *P. myrtifolia* were spiked with individual strains representing each subspecies of *Xf* (Xff: CFBP 7970, *Xfmo*: CFBP 8084, *Xfp:* CFBP 8402, *Xfm*: CFBP 8416). Other plant materials were spiked with the strain representing the only subspecies that infects them naturally. However, as several subspecies may co-occur in a same area and plant species may be hosts of several subspecies, samples of *N. oleander*, *O. europaea*, *P. dulcis*, and *P. myrtifolia* were also spiked with duos or trios of strains. A total of 29 plant species - *Xf* subspecies were combined. For negative controls, the samples were directly ground in sterile distilled water (5 mL). Samples were treated as above before DNA extraction. All DNA extractions were performed using the QuickPickTM SML Plant DNA Kit (Bio-Nobile, Turku, Finland) as in PM7/24 (EPPO, 2018b) with an automated system (Caliper Zephyr, PerkinElmer). A control composed of DNAs extracted from bacterial suspensions were systematically performed.

### 2.5 Relationships between DNA concentration, OD_600nm_ and bacterial concentration

Fresh suspensions of CFBP 7970 strain calibrated at OD_600nm_ = 0.1 were plated on PWG medium and incubated at 28°C for 8 days before counting. They contained 1×10^8^ CFU.mL^−1^. Genomic DNA from the same suspensions was extracted using QuickPickTM SML Plant DNA Kit (Bio-Nobile, Turku, Finland) as described in PM7/24 (EPPO, 2018b). DNA concentration was measured using Qubit fluorimeter and serial dilutions of *Xf* genomic DNA at concentrations ranging from 1 µg.mL^−1^ to 1 pg.mL^−1^ were prepared. The DNA was amplified using the Harper’s *et al*. (2010) qPCR assay in a Bio-Rad CFX384 thermocycler. Results of the amplified serial dilutions were used to establish standard curves relating the amount of fluorescence to the amount of DNA. The bacterial concentration of the corresponding DNA solution was calculated based on DNA measures using an estimated genome size of 2,493,794 bp for the strain CFBP 7970 (Denancé *et al*., 2017) and knowing that 1 pg = 9.78×10^8^ bp (Doležel *et al*., 2003). Using the following equation curve (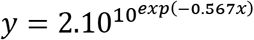, R^2^ = 0.999) a Ct = 19.8 correlated to 1.04 × 10^8^ genome equivalent.mL^−1^.

### 2.6 Gene target selection and primers design

SkIf tool (Briand *et al*., 2016) was used on 58 *Xylella* genomic sequences to target specific sequences of the *Xf* species, each subspecies, and the *fastidiosa sensu lato* (*Xffsl*) subspecies, i.e. the group including the *fastidiosa*, *morus* and *sandyi* subspecies (Denancé *et al*., 2019) (Table 2). Six primer and probe combinations were designed using Primer3 2.3.4 (Koressaar and Remm, 2007), on these specific sequences to target the whole *Xf* species (XF primers), and the various subspecies: *fastidiosa* (XFF primers), *fastidiosa sensu lato* (XFFSL primers), *morus* (XMO primers), *multiplex* (XFM primers) and *pauca* (XFP primers) (Table 3). The parameters were set up with an optimal size of 20 bp (sizing between 18-27 bp), an optimal product size of 85 to 150 bp; a Tm of 60°C (± 3°C) and 70°C (± 3°C) for primers and probes, respectively. Then, the individual primer and probe combinations and the six sets of four combinations were tested using Amplify to check the absence of dimer and cross-amplification (Engels, 1993). The specificity of all primers and probes was tested *in silico* using PrimerSearch (Val Curwen, Human Genome Mapping Project, Cambridge, UK) on the initial set of 58 genomic sequences of *Xylella* and on the 154,478 bacterial Whole Genome Shotgun (WGS) sequences available in the NCBI database (as on August 22, 2018). BLASTn of the amplicons were run on the NCBI WGS database to evidence their specificity.

**Table 2:** Description and composition of the longest specific long-mers obtained using SkIf for the various targets.

| Target <sup>a</sup> | Long-mer size (bp) | Long-mer position (in the genome of the given strain) | Targeted CDS: locus name, position | Putative function |
| --- | --- | --- | --- | --- |
| **XF** | 986 | 1,254,689 - 1,255,674 (M23) | WP_004084873,<br>1,254,698 - 1,255,674 | Ketol-acid reductoisomerase |
| **XFF** | 516 | 2,477,123 - 2,477,638 (M23) | ACB93575,<br>2,476,428 - 2,477,645 | Restriction modification system |
| **XFFSL** | 227 | 719,367-719,593 (M23) | ACB92051,<br>719,717 - 718,980 | Unknown |
| **XFM** | 1660 | 1,825,046-1,826,705 <sup>b</sup> (M12) | WP_004083558,<br>1,824,865 -1,825,101 | Unknown |
|  |  |  | WP_004083559,<br>1,825,613 - 1,825,855 / | Unknown |
|  |  |  | WP_004083560,<br>1,826,106 - 1,826,489 / | DNA adenine methylase |
|  |  |  | WP_004083562,<br>1,826,593 - 1,826,768 | DNA adenine methylase |
| **XFMO** | 288 | 1,908,250-1,908,548 (MUL0034) | AIC14009,<br>1,908,261 - 1,908,798 | Peptidase S24 |
| **XFP** | 876 | 337,676 - 338,551 <sup>b</sup> (De Donno) | ARO67912,<br>336,864 - 338,246 / | Histidine kinase |
|  |  |  | ARO69620,<br>338,246 - 339,286 | ABC transporter substrate-binding |

<sup>a</sup>: see Table 3 for description of codes of primer and probe sets
<sup>b</sup>: the long-mer is overlapping several CDS

**Table 3:**
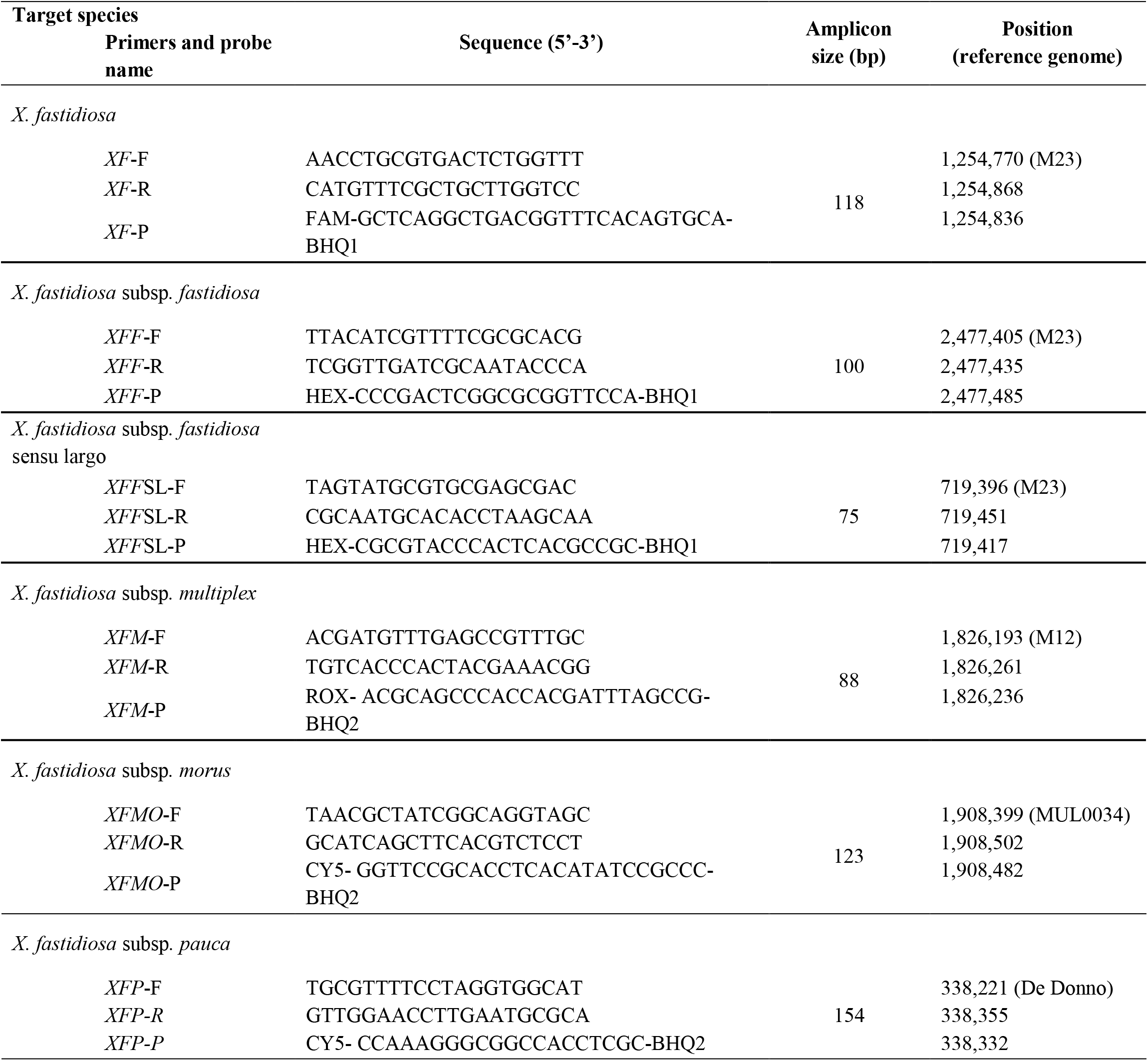
Primers and probes designed in this study for *Xf* detection at the species and subspecies level.

Four others primer and probe combinations previously published were used in this study. The first targets the *rimM* gene of *Xf* (Harper *et al*., 2010) and was used as reference protocol. The second targets the eukaryotic *rRNA18S* gene (Ioos *et al*., 2012) and was used as internal control. The remaining two tests target *fastidiosa* or *multiplex* subspecies (Burbank and Ortega, 2018).

### 2.7 Optimization of qPCR assays and tetraplexing

The tetraplex qPCR assays designed in this study were optimized for: i) primer and probe hybridization temperature that was checked individually by PCR using a gradient ranging from 57.5 to 61.4°C in intervals of 0.8°C (CFX96 Touch™ Bio-Rad), ii) concentrations of 250 nM, 575 nM or 900 nM for primers combined with 150 nM, 200 nM or 250 nM for probes according to PCR mix manufacturer instructions, and iii) addition of 600 ng.µl^−1^ of BSA. All the optimization analyses were performed in triplicates using SsoAdvanced™ Universal Probes Supermix (Bio-Rad) and performed in a Bio-Rad CFX thermocycler using the “all channels” reading mode. To allow simultaneous detection of *Xf* and identification at the subspecies level, primer and probe combinations were then declined in six different triplex and tetraplex qPCR sets, *i.e.* set n°1: XF-XFFSL-XFM-XFP, set n°2: XF-XFF-XFM-XFP, set n°3: XF-XFF-XFM-XMO, set n°4: XFFSL-XFM-XFP, set n°5: Harper-XFFSL-XFM-XFP and set n°6: 18S-XFFSL-XFM-XFP.

The optimized final reaction conditions were performed in a final volume of 10 µL containing 1X of SsoAdvanced™ Universal Probes Supermix (Bio-Rad), 575 nM of primers, 200 nM of probes and 600 ng.µl^−1^ of BSA (ThermoFisher) and 1 µL of extracted DNA. The optimal thermocycling conditions selected were: 3 min at 95°C, followed by 40 cycles of 15 s at 95°C and 30 s at 60°C. The qPCR assays results were analyzed, with expert verification, using Bio-Rad CFX Manager 3.1 software and its regression mode. The reaction efficiency was calculated using serial dilutions with the formula: E = 10^(−1/slope)^.

### 2.8 qPCR assay specificity, efficiency and limit of detection

The specificity of the newly designed primer and probe combinations was validated using the optimized protocol on the boiled bacterial suspensions of the 69 strains listed in the Table 1. The efficiency of each combination was evaluated on bacterial DNA solutions ranging from 1 µg.mL^−1^ to 1 pg.mL^−1^, in simplex or tetraplex assays (set n°1 to 3), on the strains CFBP 7970 (*Xff*) for the primers XF, XFF and XFFSL, CFBP 8416 (*Xfm*) for the primers XF and XFM, CFBP 8084 (*Xfmo*) for the primers XF and XFMO, and CFBP 8402 (*Xfp*) for the primers XF and XFP. In addition, each set was also evaluated with spiked plant material. All analyses were performed in triplicate. Two independent experiments were carried out on *O. europaea, P. myrtifolia, P. cerasus, P. dulcis Q. ilex* and *V. vinifera* using the set n°1: XF – XFFSL – XFM – XFP, leading to the analysis of 46 combinations of plant/strain(s) for this set. The assays were also performed on environmental plant samples and inoculated plant samples. For plant samples, the lowest concentration with a positive result in at least two out of the three replicates was considered the limit of detection (LOD).

The LOD of the tetraplex qPCR assays sets n°1 to 3 was compared to the Harper’s qPCR detection test (Harper *et al*., 2010) using the TaqMan™ Universal PCR Master Mix (Applied Biosystems) as in PM7/24 (EPPO, 2018b). The LOD of the tetraplex qPCR assay set n°1 was compared to the ones of sets n°4, 5 and 6. The specificity of the qPCR assay recently proposed by Burbank and Ortega (2018) was also evaluated on the *Xf* strain collection usingthe SsoAdvanced™ Universal Probes Supermix (Bio-Rad) master mix.

## 3 Results

### 3.1 Design of primers and probes and *in silico* analysis

Species-specific and subspecies-specific long-mers were identified with SkIf (Briand *et al*., 2016; Denancé et al., 2019) on genomic sequences. For the *Xf* species and the subspecies *fastidiosa, morus, multiplex*, and *pauca*, one of the two longest long-mers identified by Denancé *et al* (2019) was selected for this study (Table 2). For the subspecies *fastidiosa sl* specific long-mers were searched for on our 58 genome sequences of *Xf*, using the subspecies *fastidiosa, morus* and *sandyi* genomes as ingroups and the *multiplex* and *pauca* genomes as outgroups. In total, 3,345 long-mers were identified, ranging from 22 bp to 235 bp (Supplemental data 1).

Primers and probes were designed within specific long-mers (Table 3). Specific amplifications were obtained *in silico* on XF genome sequences and WGS bacterial sequences from NCBI at the expected amplification size, without any mismatch for the five primer and probe combinations (XFF, XFFSL, XFM, XFMO and XFP). Only two mismatches were observed and concerned the XF primer and probe combination. One mismatch was on the eighth nucleotide on the XF probe for the *Xfm* Dixon, Griffin1, M12, Sycamore, CFBP 8416, CFBP 8417, CFBP 8418 strains and the second one was on the sixth nucleotide of the forward XF primer of the Ann-1 *Xfs* strain. As there were not many possible combinations of primers and probes for the XF set, this combination was nevertheless retained, and subsequent *in silico* checks proved the specificity of all primer and probe combinations.

### 3.2 Specificity and sensitivity of simplex and tetraplex qPCR assays on strains

The specificity of each newly designed primer and probe combination was validated in simplex qPCR assays on 39 *Xf* strains and on 30 plant associated-bacterial strains (Table 1). These strains were selected as they potentially share the same niche than Xf or for being phylogenetically closely related. No amplification was detected on non-target strains or healthy host plant species and the primer and probe combinations allowed amplification of all strains or subspecies of *Xf*, for which they were designed (XF: 39/39, XFF: 10/10, XFM: 16/16, XFMO: 1/1, XFP: 7/7, XFFSL: 16/16).

In simplex qPCR assays, the LODs of the new primer and probe combinations designed in this study were as good as the LODs obtained with the Harper’s qPCR assay or 10 times better for XFM primers (Table 4). The efficiency of each combination was evaluated on serial dilutions of calibrated DNA solutions. The XF, XFM, XFMO, XFP, and XFFSL primers and probes allowed detection of *Xf* up to 10 pg.mL^−1^ (4 copies/reaction). XFF primers were slightly less sensitive with a threshold up to 100 pg.mL^−1^ (40 copies/reaction). On the same DNA solutions, Harper *et al*. (2010) qPCR assay allowed the detection of strains CFBP 8402 (*Xfp)* and CFBP 8084 (*Xfmo*) up to 10 pg.mL^−1^, and CFBP 7970 (*Xff*) and CFBP 8416 (*Xfm*) strain up to 100 pg.mL^−1^. This makes our new primer qPCR assays good alternatives to Harper’s qPCR assay.

**Table 4:** Efficiency of the primer and probe sets in simplex qPCR assays on extracted DNA of bacterial strains in comparison with the Harper’s test (Harper *et al.*, 2010). A, Mean Ct value for each primer and probe set on target strains; B, Percentage of efficiency and standard curve parameters of each primer and probe set on target strains.

| DNA concentration | Theoretical number of genome copy.mL<sup>-1</sup> | XF (CFBP 7970) | XFF (CFBP 7970) | XFM (CFBP 8416) | XFMO (CFBP 8084) | XFP (CFBP 8402) | XFFSL (CFBP 7970) | XFFSL (CFBP 8084) | Harper's (CFBP 7970) | Harper's (CFBP 8416) | Harper's (CFBP 8084) | Harper's (CFBP 8402) |
| --- | --- | --- | --- | --- | --- | --- | --- | --- | --- | --- | --- | --- |
| 1 µg.mL<sup>-1</sup> | 4x10<sup>8</sup> | 20.03<sup>a</sup><br>(0.08) | 18.47<br>(0.16) | 19.34<br>(0.04) | 19.09<br>(0.03) | 16.64<br>(0.12) | 18.67<br>(0.01) | 18.94<br>(0.04) | 17.82<br>(0.02) | 17.36<br>(0.05) | 17.80<br>(0.04) | 16.58<br>(0.04) |
| 100 ng.mL<sup>-1</sup> | 4x10<sup>7</sup> | 23.31<br>(0.10) | 21.88<br>(0.07) | 22.80<br>(0.10) | 22.78<br>(0.10) | 19.63<br>(0.06) | 22.09<br>(0.05) | 23.10<br>(0.08) | 21.45<br>(0.33) | 21.03<br>(0.09) | 22.13<br>(0.34) | 19.23<br>(0.03) |
| 10 ng.mL<sup>-1</sup> | 4x10<sup>6</sup> | 26.56<br>(0.03) | 25.49<br>(0.06) | 26.18<br>(0.09) | 25.91<br>(0.07) | 22.93<br>(0.10) | 26.84<br>(1.01) | 27.55<br>(0.06) | 25.88<br>(0.06) | 25.35<br>(0.12) | 25.55<br>(1.55) | 22.76<br>(0.04) |
| 1 ng.mL<sup>-1</sup> | 4x10<sup>5</sup> | 30.22<br>(0.19) | 28.65<br>(0.07) | 29.06<br>(0.12) | 28.89<br>(0.08) | 25.95<br>(0.07) | 28.61<br>(0.24) | 30.78<br>(0.04) | 29.98<br>(0.16)<sup>a</sup> | 29.02<br>(0.11) | 29.36<br>(0.11) | 25.77<br>(0.15) |
| 100 pg.mL<sup>-1</sup> | 4x10<sup>4</sup> | 33.36<br>(0.43) | 31.57<br>(0.18) | 32.42<br>(0.37) | 32.18<br>(0.20) | 28.95<br>(0.08) | 31.82<br>(0.85) | 33.44<br>(0.16) | na | na | 32.53<br>(0.20) | 31.55<br>(0.16) |
| 10 pg.mL<sup>-1</sup> | 4x10<sup>3</sup> | 36.28<br>(1.36) | na | 37.37<br>(0.72) | 36.07<br>(0.59) | 31.82<br>(0.59) | 33.86<br>(3.63) | 38.52<br>(0.08) | na | na | na | 34.28<br>(0.73) |
| 1 pg.mL<sup>-1</sup> | 4x10<sup>2</sup> | na<sup>b</sup> | na | na | na | na | na | na | na | na | na | na |

<sup>a</sup>: a signal is considered positive when obtained in at least two of the three technical repetitions and the lowest concentration at which a signal is obtained is the LOD
<sup>b</sup>: not detected

**B.**
| Target | Strain code | Efficiency | R<sup>2</sup> | Slope |
| --- | --- | --- | --- | --- |
| XF | CFBP 7970 | 101.4% | 0.978 | -3.289 |
| XFF | CFBP 7970 | 101.1% | 0.997 | -3.297 |
| XFM | CFBP 8416 | 100.4% | 0.995 | -3.311 |
| XFMO | CFBP 8084 | 100.0% | 0.996 | -3.299 |
| XFP | CFBP 8402 | 112.6% | 0.995 | -3.052 |
| XFFSL | CFBP 7970 | 95.5% | 0.996 | -3.434 |
| XFFSL | CFBP 8084 | 102.0% | 0.957 | -3.274 |

The three tetraplex qPCR assays (set n°1: XF – XFFSL – XFM – XFP, set n°2: XF – XFF – XFM – XFP and set n°3: XF – XFF – XFM – XFMO) allowed both detection and identification of *Xf* and its subspecies (Supplemental data 2). On calibrated DNA solutions these assays were as good as Harper’s test or had a LOD 10 times higher depending of the tetraplex assays. When used in tetraplex the Ct values obtained were always lower than the Ct values obtained with Harper’s test. Except for *morus* primers (XFMO) the LOD of tetraplex qPCR assays was usually 10 times higher than the LOD of the simplex test on DNA (Table 4 and Supplemental data 2). In addition, it should be noted that the closer the Ct value was to the detection limit, the higher the SEM was. In tetraplex qPCR assays set n°1, XF, XFM and XFP primers allowed a detection up to 100 pg.mL^−1^. The XFFSL primers allowed the detection of *Xff* up to 10 pg.mL^−1^ and of *Xfmo* up to 100 pg.mL^−1^. The set n°2 allowed detection up to 100 pg.mL^−1^ using XFF and XFM primers and up to 10 pg.mL^−1^with XFP primers. The XF primers allowed the detection of *Xff* and *Xfm* up to 100 pg.mL^−1^and of *Xfp* up to 10 pg.mL^−1^. The set n°3, allowed a detection up to 100 pg.mL^−1^ with XF, XFF and XFM primers and up to 10 pg.mL^−1^ with XFMO primers.

A triplex qPCR assay for the simultaneous detection of subspecies *fastidiosa* and *multiplex* has recently been published (Burbank and Ortega, 2018). In order to analyze the potential of their targets and potentially introduce them into our sets to improve *Xf* detection, we tested their specificity *in silico* and *in vitro* on selected bacterial strains. According to BLASTn searches, *Xff* primers potentially amplified two of the three strains of the subsp. *sandyi* (CFBP 8073: ST75 and Co33: ST72) without mismatches and seven strains of the subsp. *pauca* (CoDiRo, COF0407, De Donno, OLS0478, OLS0479, Salento-1 and Salento-2) with one and two mismatches on the forward and reverse primers, respectively (Supplemental data 3). *In silico, Xfm* primers potentially amplified eight strains of subsp. *pauca* (CFBP 8072, CoDiRo, COF0407, De Donno, OLS0478, OLS0479, Salento-1, Salento-2) with three mismatches on the forward primer, two mismatches on the reverse primer and one mismatch on the probe, and amplicons had the expected size. We double checked the specificity of these two sets *in vitro* on bacterial suspensions (Supplemental data 4). *Xff* primers amplified the three tested strains of subsp. *sandyi* (CFBP 8356, CFBP 8419 and CFBP 8077) and six of the seven tested strains of subsp. *pauca* (CFBP 8074, CFBP 8402, CFBP 8429, CFBP 8477, CFBP 8495 and CFBP 8498). The sequencing of all amplicons confirmed the results of the qPCR assays. *Xfm* primers amplified five of the seven tested strains of *Xf* subsp. *pauca* (CFBP 8072, CFBP 8074, CFBP 8402, CFBP 8495 and CFBP 8498). Burbank and Ortega (2018) used a cut off at Ct=35 for categorizing a result as positive. In that case only two *pauca* strains (CFBP 8072 and CFBP 8495) would have been identified as *Xfm*, the others having values ranging between 35.33 and 35.83. For *Xfm*, due to the high Ct values, no sequencing was feasible to confirm the identification.

### 3.3 Identification of *Xf* subspecies in spiked samples with tetraplex qPCR assays

After validation of the efficiency and specificity of the primers and probe, the three sets of tetraplex qPCR assays n°1, 2 and 3, were tested on spiked samples. As the three sets gave similar results, this section is focused on the tetraplex set n°1: XF – XFFSL – XFM – XFP, which covers the full known diversity of *Xf* (Table 5). The results of the other two tetraplex assays are provided in Supplemental Data 5 and Supplemental data 6. This tetraplex qPCR assay (set n°1) was tested on 29 combinations of plant petioles and midribs spiked with one to three strains of the different subspecies. (The full results of the dilution ranges are available in Supplemental data 7). This tetraplex allowed the detection and correct identification of all subspecies in all combinations without false positive result. Although the detection limit was expected to be similar for all plants, since they were all enriched with the same bacterial suspensions, different LODs were observed ranging from 1×10^3^ to 1×10^5^ CFU.mL^−1^ (5 to 5×10^3^ CFU/reaction) depending on the matrix for plants spiked with only one strain. An independent repetition of this test was performed two months after the first one. For *O. europaea, P. myrtifolia, P. cerasus, P. dulcis* and *Q. ilex* the LOD was either identical between the two assays or 10 time higher. The LOD of *Xf* in *V. vinifera* was 100 times higher in the second assay highlighting a potential accumulation of qPCR inhibitors between the two experiments. Moreover, on 11 combinations out of 46, XF primers had a LOD 10 times higher *in planta* than the one obtained for the subspecies. *Xf* subspecies could be identified until a Ct value of 35.08 using Harper’s qPCR assay in a spiked sample of *P. dulcis*. In other matrices the LOD of the tetraplex qPCR assay corresponded usually to a Ct value ranging from 30 to 34 using Harper’s qPCR.

**Table 5:**
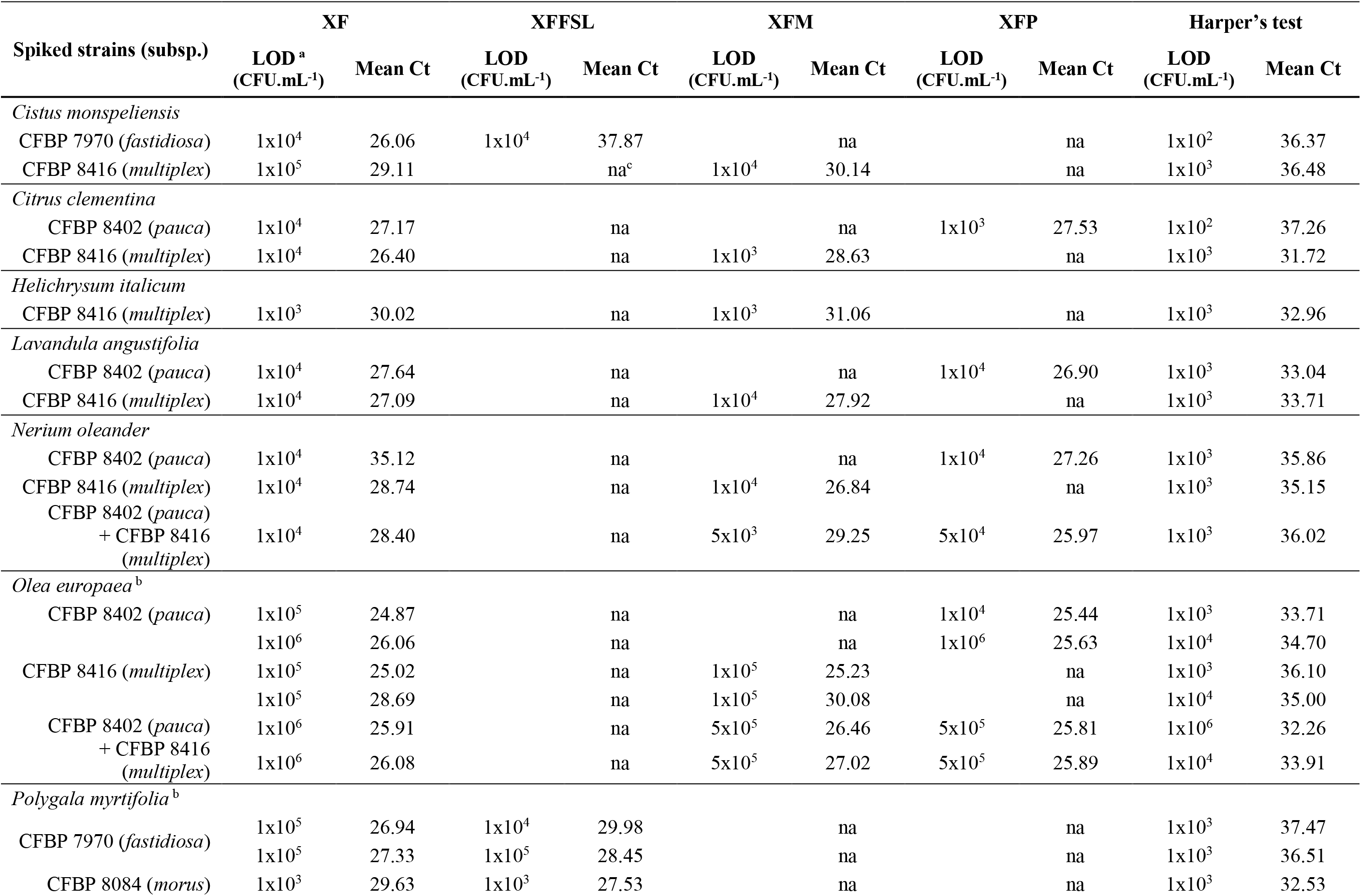
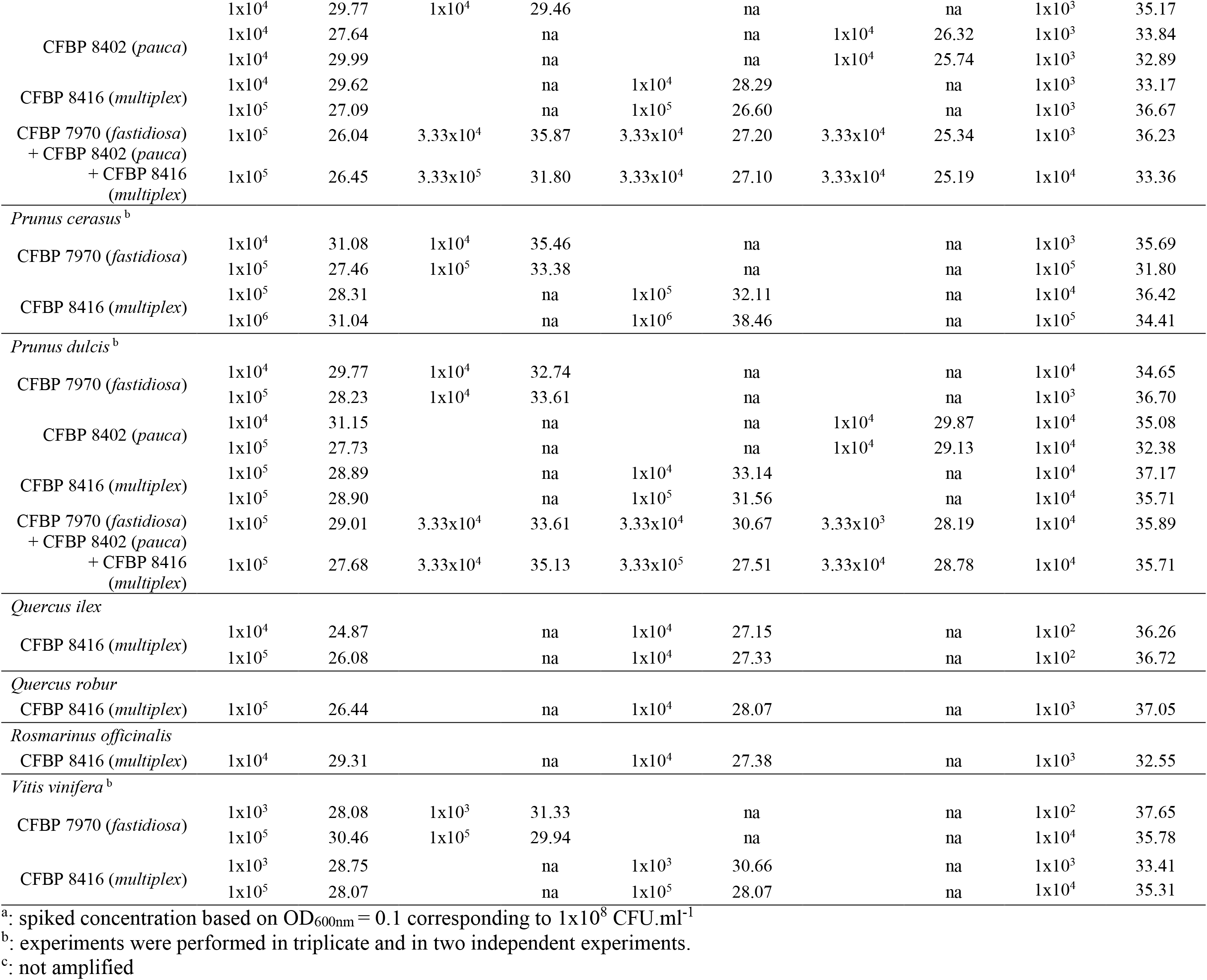
Limit of detection (LOD) of *X. fastidiosa* strains in spiked matrices using the tetraplex qPCR assay XF – XFFSL – XFM – XFP (set n°1) in comparison with the reference test (Harper’s test, Harper et al., 2010).

Moreover, the tetraplex qPCR assay set n°1 allowed the detection and identification of mix infections with two to three subspecies simultaneously. On *N. oleander, O. europaea, P. myrtifolia* and *P. dulcis* the LOD for the two or three inoculated subspecies is similar of the one obtained for single inoculations (Table 5).

To demonstrate that our multiplex qPCR assays are modular tools, which can be adapted to one’s needs, three other primer and probe sets were evaluated. In one set, we removed the primers and probe targeting the species (set n°4: XFFSL-XFM-XFP). In a second one, we replaced it by the Harper’s primers and probe as this test is known to be highly sensitive (set n°5: Harper-XFFSL-XFM-XFP), and we also tested the use of primers and probes targeting the 18S rRNA as an internal control (set n°6: 18S-XFFSL-XFM-XFP). Evaluation of these three sets on calibrated DNA suspensions of the *Xff* strain CFBP 7970 indicated that the LOD for the XFFSL primers was the same than the one found previously for the sets n°1, 4, 5 and 6 (10 pg.mL^−1^) (Supplemental data 8). In *Q. robur* and *C. monspeliensis* samples spiked with the *Xfm* strain CFBP 8416, the LOD obtained for the primers detecting the multiplex subspecies (XFM) was the same for the three sets (1×10^5^ CFU.mL^−1^) (Supplemental data 9). The use of Harper’s primers and probe in set n°5 allowed the detection of *Xf* strain at the same LOD than for XF primers and probe in spiked *Q. robur* samples, but the detection was slightly better (a gain of one Log unit) in the spiked *C. monspeliensis* samples. A Ct value was obtained for all spiked samples with the 18s rRNA primers, highlighting that these primers and probe were reliable internal amplification controls.

### 3.4 Identification of *Xf* subspecies in environmental plant samples and inoculated plants by tetraplex qPCR assays

Ten plant samples from Corsica, France (Table 6) and ten samples from inoculated plants (Table 7) were tested using the tetraplex set n°1. Our tetraplex qPCR assay was able to detect the bacterium in samples declared contaminated with Harper’s qPCR assay up to Ct =34.97. However, this LOD was variable depending on the matrices (Table 7). While the bacterium was detected at the subspecies level with one or the other primer and probe combinations in eight environmental plant samples, the XF primers and probe was less efficient and allowed the detection in only five samples (Table 6) indicating that primer and probe combinations designed for subspecies were more sensitive than the one designed to detect the species. The subspecies was hence identified in samples that were not successfully typed using the MLST protocol. Samples of *Centranthus trinervis, Olea europaea* and *Phylirea angustifolia* (n° 1, 6 and 7) were infected by a *Xffsl* strain and samples of *Helichrysum italicum*, *Lavandula stoechas*, *Polygala myrtifolia*, and *Spartium junceum* (n°2, 3, 8, 9 and 10) were detected infected by a *multiplex* strain. The partial MLST subspecies identification of the sample n°8 was hence validated. The assay also identified the subspecies in the ten samples obtained from inoculated plants and confirmed the identity of the inoculated strain.

**Table 6:** Detection of *X. fastidiosa* in environmental plant samples with low population sizes using the tetraplex qPCR assay set n° 1 in comparison with the reference test (Harper’s test, Harper et al., 2010).

| Sample | Host plant | Place (year) | Mean Ct (SEM)<sup>a</sup> |  |  |  |  | typing |
| --- | --- | --- | --- | --- | --- | --- | --- | --- |
|  |  |  | XF | XFFSL | XFM | XFP | Harper's test |  |
| 1 | *Centranthus trinervis* | Bonifaccio, France (2017) | na<sup>b</sup> | 33.67 (1.42) | na | na | 34.97 (0.53) | unknown |
| 2 | *Helichrysum italicum* | Propriano, France (2017) | 27.35 (0.67) | na | 27.25 (0.23) | na | 30.85 (0.04) | unknown |
| 3 | *Lavandula stoechas* | Vignola, France (2017) | 30.75 (0.73) | na | 26.27 (0.38) | na | 29.50 (0.13) | unknown |
| 4 | *Lavandula stoechas* | Propriano, France (2017) | na | na | na | na | 34.81 (1.40) | unknown |
| 5 | *Olea europaea* | Afa, France (2017) | na | na | na | na | 34.01 (0.77) | unknown |
| 6 | *Olea europaea* | Vignola, France (2017) | na | 29.91 (0.80) | na | na | 32.94 (0.18) | unknown |
| 7 | *Phyllirea angustifolia* | Bonifaccio, France (2017) | na | 30.52 (0.21) | na | na | 33.99 (1.09) | unknown |
| 8 | *Polygala myrtifolia* | Vignola, France (2017) | 24.86 (0.04) | na | 25.00 (0.03) | na | 25.96 (0.04) | suspected *Xfm*<sup>c</sup><br>leuA: 3 |
| 9 | *Polygala myrtifolia* | Porto-Vecchio, France (2018) | 30.14 (0.58) | na | 29.52 (0.17) | na | 32.82 (0.41) | unknown |
| 10 | *Spartium junceum* | Corbara, France (2017) | 23.68 (0.17) | na | 23.97 (0.14) | na | 24.97 (0.06) | unknown |

<sup>a</sup>: none of these test was performed by the French national reference laboratory
<sup>b</sup>:not amplified
<sup>c</sup>: typing is suspected when the seven housekeeping genes could not be amplified

**Table 7:**
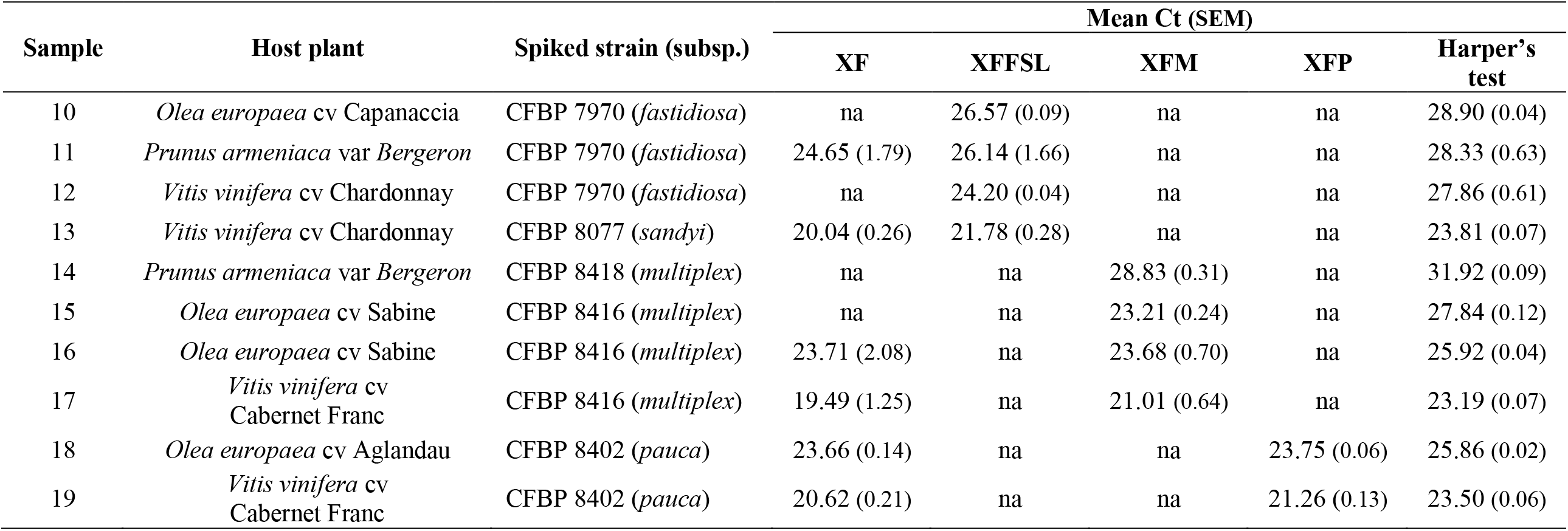
Detection of *X. fastidiosa* in inoculated plants using the tetraplex qPCR assay (set n° 1) in comparison with the reference test (Harper’s test, Harper et al., 2010).

| Sample | Host plant | Spiked strain (subsp.) | Mean Ct (SEM) |  |  |  |  |
| --- | --- | --- | --- | --- | --- | --- | --- |
|  |  |  | XF | XFFSL | XFM | XFP | Harper's test |
| 10 | *Olea europaea* cv Capanaccia | CFBP 7970 (*fastidiosa*) | na | 26.57 (0.09) | na | na | 28.90 (0.04) |
| 11 | *Prunus armeniaca* var Bergeron | CFBP 7970 (*fastidiosa*) | 24.65 (1.79) | 26.14 (1.66) | na | na | 28.33 (0.63) |
| 12 | *Vitis vinifera* cv Chardonnay | CFBP 7970 (*fastidiosa*) | na | 24.20 (0.04) | na | na | 27.86 (0.61) |
| 13 | *Vitis vinifera* cv Chardonnay | CFBP 8077 (*sandyi*) | 20.04 (0.26) | 21.78 (0.28) | na | na | 23.81 (0.07) |
| 14 | *Prunus armeniaca* var Bergeron | CFBP 8418 (*multiplex*) | na | na | 28.83 (0.31) | na | 31.92 (0.09) |
| 15 | *Olea europaea* cv Sabine | CFBP 8416 (*multiplex*) | na | na | 23.21 (0.24) | na | 27.84 (0.12) |
| 16 | *Olea europaea* cv Sabine | CFBP 8416 (*multiplex*) | 23.71 (2.08) | na | 23.68 (0.70) | na | 25.92 (0.04) |
| 17 | *Vitis vinifera* cv Cabernet Franc | CFBP 8416 (*multiplex*) | 19.49 (1.25) | na | 21.01 (0.64) | na | 23.19 (0.07) |
| 18 | *Olea europaea* cv Aglandau | CFBP 8402 (*pauca*) | 23.66 (0.14) | na | na | 23.75 (0.06) | 25.86 (0.02) |
| 19 | *Vitis vinifera* cv<br>Cabernet Franc | CFBP 8402 (*pauca*) | 20.62 (0.21) | na | na | 21.26 (0.13) | 23.50 (0.06) |

## 4 Discussion

Since its first detection in Europe in 2013, *Xf* has been reported in various EU member states and on a wide host range (https://ec.europa.eu/food/sites/food/files/plant/docs/ph_biosec_legis_emergency_db-host-plants_update12.pdf). It is hence considered as an emergent plant bacterium in Europe and it is regulated in the EU as a quarantine organism under Council Directive 2000/29/EC. Control measures to prevent the spread of this pathogen within the EU are limited to eradication and containment measures (EFSA, 2018b). Application of these outbreak management strategies require the identification of *Xf* strains at the subspecies level. Indeed, the list of host plants is provided per *Xf* subspecies with only a limited number of plants (currently 15) being hosts of all subspecies currently detected in the EU. Identifying *Xf* at the subspecies level is thus highly important to limit the number of host plants to be eradicated once an outbreak is detected.

In this context, on the basis of a large dataset of in-house and publicly available genome sequences of *Xf* and SkIf, a powerful bioinformatics-tool (Briand *et al*., 2016; Denancé *et al*., 2019), we identified species and subspecies signatures. These long-mers were used as targets to designed primer and probe combinations with different levels of specificity. These combinations target single-copy genes encoding proteins involved in bacterial metabolism. This is the case for the XF primers and probe targeting a gene encoding a ketol-acid reductoisomerase, an enzyme essential in the biosynthesis pathway of the L-isoleucine and L-valine; XFF primers and probe target a gene encoding a restriction modification system DNA specificity, involved in defense against foreign DNA (Wilson and Murray, 1991); XFM primers and probe target a gene coding a DNA methyltransferase; XFMO primers and probe target a gene coding an S24 peptidase involved in a stress-response against DNA lesions and leading to the repair of single-stranded DNA (Erill *et al*., 2007); XFP primers and probe target a gene coding a histidine kinase and an ABC transporter substrate, two membrane proteins involved in signal transduction across the cellular membrane (Yoshida *et al*., 2007; Tanaka *et al*., 2018). The targets of our subspecific assays were selected to be exactly identical among all strains of a given subspecies and absent from any other bacteria, thus these targets are not recombining elements.

Tested on a large collection of target and non-target strains, the primers and probes showed high specificity for *Xf* and its subspecies and no cross-reactions. *In vitro*, the specificity was tested in two steps. Inclusivity was evaluated on strains of *Xf* subspecies and exclusivity on a range of strains chosen to be present in the same plant and insect niches as *Xf* (Rogers, 2016) or to be genetically closely related to it. With the exception of a few studies (Boureau *et al*., 2013; Hulley *et al*., 2019) only one to ten non-target strains are selected to test the specificity of novel molecular detection tools (Francis *et al*., 2006; Harper *et al*., 2010; Burbank and Ortega, 2018). Here a larger collection including 30 non-target strains and 39 *Xf* strains was analyzed to ensure the specificity of the primer and probe combinations based on the advice of the PM 7/98 of the EPPO (2014) and the MIQE of Bustin *et al*. (2009).

At the moment there is only few methods allowing the simultaneous detection and identification of different subspecies of *Xf* and none of them is specific. The conventional PCR test of Hernandez-Martinez *et al*. (2006) was designed to differentiate the subspecies *multiplex, fastidiosa* and *sandyi*. Nevertheless, the analysis of more than 300 samples collected in France and infected with subsp. *multiplex* revealed the amplification of additional bands leading to unclear patterns (Denancé *et al*., 2017). A triplex qPCR assay was recently developed to identify *Xff* and *Xfm* and was tested on grapevine, almond and insects (Burbank and Ortega, 2018). Compared to this assay, our tetraplex qPCR assays gave similar results for the analysis of spiked almond and grapevine samples. However, we did not detect any cross reaction with our primers and probes, while the test proposed by Burbank and Ortega in 2018 could lead to cross-reactions with strains from the subspecies *pauca* and *sandyi.* While *pauca* strains have not been so far detected in grapevine samples in any outbreaks, it was demonstrated that grapevine is susceptible to *pauca* strains (Li et al., 2013) and caution should be taken not to misidentify *Xf* strains infecting grapevine.

Primers and probes optimized for qPCR tetraplex assays allowed simultaneously the detection of *Xf* and its identification at the subspecies level, providing two complementary results as the targets of the tests are different. The use of one of these tetraplex assays hence corresponds to the first requirement for *Xf* detection as reported in PM 7/98 (EPPO, 2014). So far, subspecies identification is done by sequencing two to seven housekeeping genes (Yuan *et al*., 2010; EPPO, 2018b). If one of the gene amplifications fails, or if sequencing is not feasible (in case of a too low amount of DNA) then the subspecies cannot be assigned. The average value of the LOD for every gene in the *Xf* MLST scheme is at the best at 10^5^ CFU.mL^−1^ (Cesbron *et al*, in prep). As demonstrated with single strain suspensions and mix-suspensions these assays display high efficiency (i.e. low LOD), even if, as Ito and Suzaki (2017) have shown, multiplexing increases the LOD by up to one log unit. With a LOD of 10 to 100 pg.mL^−1^ (i.e. 4×10^3^ to 4×10^4^ copies.mL^−1^), these multiplex qPCR assays still present a sensitivity that is similar to the one of the reference protocol, on single bacterial suspensions (Harper *et al*., 2010).

In spiked and environmental plant samples, the benefit from the use of our tetraplex assays is obvious. The tetraplex qPCR assays developed here are able to identify *Xf* subspecies up to 10^3^ CFU.mL^−1^ in spiked samples. They allowed the identification of the *Xf* subspecies in environmental plant samples, as well, leading to subspecies identification when MLST failed and confirmed partial MLST identification. Subspecies was identified in samples detected infected but with high Ct values (determined at 35 with the Harper’s qPCR assay), which corresponds to a bacterial load of only 10^3^ CFU.mL^−1^. It should be mentioned here, that to increase the chance of detecting *Xf* in low contaminated samples, a sonication step has been added before DNA extraction. Indeed, it has been known for a while that sonication allows bacterial recovery from plant samples (Morris *et al*., 1998) and this was recently demonstrated to improve *Xf* isolation from plant samples (Bergsma-Vlami *et al*., 2017). We hypothesize that a sonication step while disrupting biofilm, will allow a better cell lysis through a better access of chemicals to the cells. Although analysis of more samples is necessary to confirm this LOD, the tetraplex qPCR assays allow the identification of *Xf* subspecies in samples for which it was not possible with the current MLST scheme, even considering only two genes.

In spiked plant samples the LOD of our tetraplex qPCR assays were 10 to 100 times higher than in bacterial suspensions. This could be linked to the presence of plant metabolites, mostly polyphenols, polysaccharides but also pectin or xylan, that act as inhibitors of the polymerase. To avoid such a problem, we already included BSA in the PCR reaction mix to chelate polyphenols (Harper *et al*., 2010; Wei *et al*., 2008). Moreover, we used polymerases that are known to be less susceptible to inhibitors than regular ones. The TaqMan™ Universal PCR Master Mix (used in the qPCR Harper’s test) contains an AmpliTaq Gold DNA polymerase, and the SsoAdvanced™ Universal Probes Supermix (Bio-Rad) (used in our tetraplex qPCR assays) contains a Sso7d fusion polymerase. Both Taq polymerases were highlighted to have good amplification performance in comparison to nine other Taq polymerases (Witte *et al*., 2018). The Sso7d fusion polymerase was optimized for multiplex qPCR and to amplify samples rich in inhibitors such as polysaccharides, cellulose or pectin. Grapevine and olive tree are known to be rich in polyphenols (Ortega-Garcia *et al*., 2008; Schneider *et al*., 2008). These compounds are accumulated in the plant during stress or fruit ripening (Ortega-Garcia *et al*., 2008; Ennajeh *et al*., 2009). These variations could explain the 10 to 100 fold higher LOD obtain for the second repetition that was performed with grapevine and olive tree sampled two months after the first sample set.

While we added a sonication step to improve DNA extraction, we did not test here other ways to improve *per se* the DNA extraction step and improve the LOD of our assays. Various options are available. A phenol-chloroform step could be added to the DNA extraction method to reduce the level of extracted proteins (Schrader *et al*., 2012). Reagents such as Tween 20, DMSO, polyethylene glycol or active carbon could be used to precipitate the polysaccharides before DNA precipitation (Schrader *et al*., 2012). Phenol levels may be reduced with the use of polyvinyl-pyrrolidone or the addition of borate (Wilkins and Smart, 1996). Drying plant samples at 65°C for 2 days, prior to DNA extraction, could also help to cancel out the effect of phenolic inhibitors (Sipahioglu *et al*., 2006).

One of the great advantages of the multiplex qPCR assays we developed is that they are modular and reliable. Combinations of primers and probe can be adapted to include sets aiming at detecting infections at the species and/or only at the subspecies level, and having internal controls for each reaction. We showed here as proofs of concept, that replacing our XF primers and probe with the ones from Harper’s test is feasible and leads to highly susceptible test, as using 18S rRNA primers and probe as internal control is efficient.

In addition, unlike with identification relying on MLST scheme, the qPCR tetraplex assays allow the simultaneous identification of several subspecies in one sample, as demonstrated with spiked samples. In fact, mix infections with two subspecies of *Xf* have already been observed in naturally infected plants (Chen *et al*., 2005; Bergsma-Vlami *et al*., 2017; Denancé *et al*., 2017). This leads to the observation of multiple peaks on the sequencing sequence of a housekeeping gene and is complex to analyze and differentiate from a sequencing error (Denancé *et al*., 2017). The simultaneous detection and identification of multiple subspecies brings the tetraplex qPCR assays powerful tools to easily and quickly detect mixed infection or to study *Xf* in areas such as Europe where several subspecies live in sympatry (Denancé *et al*., 2017).

When a new assay is developed, the time and cost difference with current protocols must be taken into account. The tetraplex qPCR assays are much faster and cheaper than using a test for detection and then a reduced MLST scheme for subspecies assignation. The current protocol costs are for Harper’s qPCR detection at the writing time ~0.52€ for reagents, (for a volume of 10 µL) ~1.62€ for the amplification of two housekeeping genes (~0.81€/gene for a volume of 20 µL) and ~10.2€ for their sequencing (~5.1€/gene in both directions), hence totalizing ~12.35€ per sample. In comparison a single tetraplex qPCR assay costs ~0.37€ per sample (for a volume of 10 µL). None of these costs includes the cost of plastic materials or specialized equipment such as a qPCR thermocycler.

To conclude, we developed specific, effective, fast, cost-efficient and easy to set up tools allowing in one step to detect and identify at the subspecies level *Xf* infection directly in plant samples. Compared to current protocols, the LOD of our tetraplex assays allowed subspecies identification at levels where regular amplifications such as the one used for MLST failed. Tetraplex qPCR assays are also easily to perform in a routine lab and as such should be easily transferable to laboratories and are modular according to the user’s needs.

## 5 Nomenclature

BLAST: Basic Local Alignment Search Tool
CNBC: National Botanical Conservatory of Corsica
INRA: French National Institute for Agricultural Research
IRHS: Research Institute of Horticulture and Seeds
LAMP: Loop-Mediated Isothermal Amplification
MIQE: Minimum Information for the Publication of Quantitative Real-Time PCR Experiments
MLST: Multi-Locus Sequence Typing
NCBI: National Center for Biotechnology Information
ST: Sequence Type
*Xf*: *Xylella fastidiosa*
*Xff*: *Xylella fastidiosa* subsp. *fastidiosa*
*Xffsl*: *Xylella fastidiosa* subsp. *fastidiosa* sensu lato
*Xfm*: *Xylella fastidiosa* subsp. *multiplex*
*Xfmo*: *Xylella fastidiosa* subsp. *morus*
*Xfp*: *Xylella fastidiosa* subsp. *pauca*
*Xfs*: *Xylella fastidiosa* subsp. *sandyi*
WGS: Whole Genome Shotgun

## 6 Acknowledgments

We thank Muriel Bahut (ANAN technical facility, SFR QUASAV, Angers, FR) for DNA extraction automatization, CIRM-CFBP (Beaucouzé, INRA, France; http://www6.inra.fr/cirm_eng/CFBP-Plant-Associated-Bacteria) for strain preservation and supply, Leonardo de la Fuente (Auburn University, AL, USA) and LSV-ANSES for sharing strains, and colleagues from CNBC for sampling plants in Corsica, France. We acknowledge Nicolas Denancé for preliminary experiments to design specific PCR tests. We thank Charles Manceau (Anses, Angers, FR) for his contribution while applying for funding, Armelle Darrasse and Matthieu Barret for fruitful discussions and critical reading of the manuscript.

## 7 Author Contributions

ED performed the experiments, ED and SC conducted the study, MB designed the bioinformatics tool, ED, MB, MAJ and SC designed the *in silico* analysis, and interpreted the data, MAJ conceived the study, and applied for funding, ED, MAJ and SC wrote the manuscript. All authors read and approved the final version of the manuscript.

## Additional files

Supplemental data 1: *Xffsl* specific kmer identified

Supplemental data 2: Efficiency of primers and probes sets multiplexed in tetraplex qPCR assays N° 1, 2 and 3 on Xf strains.

Supplemental data 3: In silico assessment of the specificity of X. fastidiosa subsp. fastidiosa (Xff) and X. fastidiosa subsp. multiplex (Xfm) primers and probe sets proposed by Burbank et al., 2018.

Supplemental data 4: Assessment of target specificity of Burbank et al., 2018 Xff and Xfm primers and probe sets using collections of strains.

Supplemental data 5: LOD of *X. fastidiosa* in spiked matrices using the tetraplex qPCR assay XF – XFF – XFM – XFP (set n°2).

Supplemental data 6: LOD of *X. fastidiosa* in spiked matrices using the tetraplex qPCR assay XF – XFF – XFM – XFMO (set n°3).

Supplemental data 7: Detection *X. fastidiosa* in dilution ranges of spiked matrices using the tetraplex qPCR assay XF – XFFSL – XFM – XFP (set n°1)

Supplemental data 8: Comparison of LOD of *X. fastidiosa* subsp. *fastidiosa* strain CFBP 7970 using the multiplex sets n°1, n°4, n°5 and n°6.

Supplemental data 9: Comparison of LOD of *X. fastidiosa* in spiked matrices using the multiplex set n°1, n°4, n°5 and n°6.

