## Supplemental data 2 to 9 for "Novel tetraplex qPCR assays for simultaneous detection and identification of *Xylella fastidiosa* subspecies in plant tissues"

### Supplemental data 1: List of *Xffsl* specific kmer identified using SkIF on genome sequences

**Xlsx file**

### Supplemental data 2: Efficiency of primer and probe sets multiplexed in tetraplex qPCR assays N° 1, 2 and 3 on [*Xf*](#_Toc8906501) strains.

| Tetraplex qPCR assay | Strain code | LOD | Theoretical number of copies.mL^-1^ | Mean Ct  (SEM) | Efficiency | R² | Slope |
| --- | --- | --- | --- | --- | --- | --- | --- |
| *Primers/Probe* |  |  |  |  |  |  |  |
| Set n°1  XF – XFFSL – XFM – XFP |  |  |  |  |  |  |  |
| *Xf* | CFBP 7970 | 100 pg.mL^-1^ | 4x10^4^ | 27.79 (0.03) | 117.5% | 0.995 | -2.963 |
| *Xf* | CFBP 8416 | 100 pg.mL^-1^ | 4x10^4^ | 28.10 (1.15) | 110.8% | 0.995 | -2.963 |
| *Xf* | CFBP 8402 | 100 pg.mL^-1^ | 4x10^4^ | 27.88 (0.12) | 106.0% | 0.990 | -3.185 |
| *Xf* | CFBP 8084 | 100 pg.mL^-1^ | 4x10^4^ | 32.17 (1.50) | 105.8% | 0.976 | -3.191 |
| *Xffsl* | CFBP 7970 | 10 pg.mL^-1^ | 4x10^3^ | 31.81 (0.13) | 120.4% | 0.997 | -2.914 |
| *Xffsl* | CFBP 8084 | 100 pg.mL^-1^ | 4x10^4^ | 34.60 (1.87) | 97.4% | 0.989 | -3.387 |
| *Xfm* | CFBP 8416 | 100 pg.mL^-1^ | 4x10^4^ | 28.78 (0.38) | 128.5% | 0.997 | -2.787 |
| *Xfp* | CFBP 8402 | 100 pg.mL^-1^ | 4x10^4^ | 26.83 (0.02) | 112.7% | 0.980 | -3.051 |
| Set n°2 XF – XFF – XFM – XFP |  |  |  |  |  |  |  |
| *Xf* | CFBP 7970 | 100 pg.mL^-1^ | 4x10^4^ | 36.05 (2.01) | 91.7% | 0.994 | -3.538 |
| *Xf* | CFBP 8416 | 100 pg.mL^-1^ | 4x10^4^ | 29.30 (0.93) | 107.2% | 0.989 | -3.160 |
| *Xf* | CFBP 8402 | 10 pg.mL^-1^ | 4x10^3^ | 34.93 (2.00) | 108.0% | 0.994 | -3.136 |
| *Xff* | CFBP 7970 | 100 pg.mL^-1^ | 4x10^4^ | 29.90 (0.20) | 97.3% | 0.994 | -3.389 |
| *Xfm* | CFBP 8416 | 100 pg.mL^-1^ | 4x10^4^ | 30.11 (0.22) | 117.1% | 0.989 | -2.971 |
| *Xfp* | CFBP 8402 | 10 pg.mL^-1^ | 4x10^3^ | 28.69 (0.16) | 109.9% | 0.989 | -3.105 |
| Set n°3 XF – XFF – XFM – XFMO |  |  |  |  |  |  |  |
| *Xf* | CFBP 7970 | 100 pg.mL^-1^ | 4x10^4^ | 30.88 (1.31) | 95.8% | 0.985 | -3.427 |
| *Xf* | CFBP 8416 | 100 pg.mL^-1^ | 4x10^4^ | 28.69 (0.65) | 109.9% | 0.994 | -3.106 |
| *Xf* | CFBP 8084 | 100 pg.mL^-1^ | 4x10^4^ | 29.75 (0.62) | 119.8% | 0.983 | -2.924 |
| *Xff* | CFBP 7970 | 100 pg.mL^-1^ | 4x10^4^ | 29.71 (0.09) | 111.8% | 0.979 | -3.069 |
| *Xfm* | CFBP 8416 | 100 pg.mL^-1^ | 4x10^4^ | 29.74 (0.44) | 117.3% | 0.995 | -2.966 |
| *Xfmo* | CFBP 8084 | 10 pg.mL^-1^ | 4x10^3^ | 29.88 (0.05) | 117.8% | 0.982 | -2.958 |

### Supplemental data 3: In silico assessment of the specificity of *X. fastidiosa* subsp. *fastidiosa* (*Xff*) and *X. fastidiosa* subsp. *multiplex* (*Xfm*) primer and probe sets proposed by Burbank et al., 2018.

| Primers/  Probe | Accession n° | Strain code | Sequence Name | Binding site | | | Nb  Mismatch  forward | Nb  mismatch  reverse | Nb  Mismatch  probe | Length  Amplified  sequence |
| --- | --- | --- | --- | --- | --- | --- | --- | --- | --- | --- |
|  |  |  |  | Primer  forward | Primer  reverse | Probe |  |  |  |  |
| *Xff* | SAMN04075659 GCA 001469395.1 | CFBP8073^a^ | LKES01000009.1 | 4510 | 80616 | 4547 | 0 | 0 | 0 | 97 |
| *Xff* | SAMN03247589 GCA 000811965.1 | CoDiRO**^b^** | JUJW01000001.1 | 10655 | 667868 | 10692 | 1 | 0 | 2 | 97 |
| *Xff* | SAMN03862123 GCA 001549735.1 | OLS0479 | LRVH01000140.1 | 3003 | 9562 | 9599 | 1 | 0 | 2 | 97 |
| *Xff* | SAMN03862124 GCA 001549755.1 | OLS0478 | LRVI01000007.1 | 9649 | 381430 | 9686 | 1 | 0 | 2 | 97 |
| *Xff* | SAMN03862125 GCA 001549825.1 | COF0407 | LRVJ01000090.1 | 9576 | 49911 | 9613 | 1 | 0 | 2 | 97 |
| *Xff* | SAMN05429913 GCA 002954185.1 | Salento-1 | CP016608.1 | 660176 | 1847826 | 1847863 | 1 | 0 | 2 | 97 |
| *Xff* | SAMN05429914 GCA 002954205.1 | Salento-2 | CP016610.1 | 660212 | 1847989 | 1848026 | 1 | 0 | 2 | 97 |
| *Xff* | SAMN06765826 GCA 002117875.1 | De Donno | CP020870.1 | 660233 | 1848137 | 1848174 | 1 | 0 | 2 | 97 |
| *Xfm* | SAMN03247589 GCA 000811965.1 | CoDiRO | JUJW01000003.1 | 64846 | 159433 | 64876 | 3 | 1 | 2 | 99 |
| *Xfm* | SAMN03862123 GCA 001549735.1 | OLS0479 | LRVH01000172.1 | 45960 | 145433 | 45990 | 3 | 1 | 2 | 99 |
| *Xfm* | SAMN03862124 GCA 001549755.1 | OLS0478 | LRVI01000021.1 | 45410 | 145544 | 45440 | 3 | 1 | 2 | 99 |
| *Xfm* | SAMN03862125 GCA 001549825.1 | COF0407 | LRVJ01000146.1 | 36187 | 43443 | 36217 | 3 | 1 | 2 | 99 |
| *Xfm* | SAMN04075656 GCA 001469345.1 | CFBP8072 | LKDK01000002.1 | 43795 | 146316 | 43825 | 3 | 1 | 2 | 99 |
| *Xfm* | SAMN05429913 GCA 002954185.1 | Salento-1 | CP016608.1 | 1805046 | 702954 | 1805076 | 3 | 1 | 2 | 99 |
| *Xfm* | SAMN05429914 GCA 002954205.1 | Salento-2 | CP016610.1 | 1805211 | 702988 | 1805241 | 3 | 1 | 2 | 99 |
| *Xfm* | SAMN06765826 GCA 002117875.1 | De Donno | CP020870.1 | 1805195 | 703173 | 1805225 | 3 | 1 | 2 | 99 |

^a^: *Xylella fastidiosa* subsp. *sandyi*

^b^: CoDiRO and all the strains in the following lines are from subsp. *pauca*

### Supplemental data 4: Assessment of target specificity of *Xff* and *Xfm* primers and probe sets proposed by Burbank *et* *al*. (2018) using our collections of strains.

| **Strain code** | **Strain subspecies** | **Mean Ct value (SEM)^a^** | |
| --- | --- | --- | --- |
|  |  | *Xff* set | *Xfm* set |
| CFBP 8072 | *pauca* | na^b^ | 32.69 (0.20) |
| CFBP 8074 | *pauca* | 33.28 (1.14) | 35.33 (0.44) |
| CFBP 8402 | *pauca* | 20.86 (0.38) | 35.83 (1.54) |
| CFBP 8429 | *pauca* | 25.65 (0.26) | na |
| CFBP 8477 | *pauca* | 21.89 (0.13) | na |
| CFBP 8495 | *pauca* | 19.87 (0.21) | 34.67 (1.66) |
| CFBP 8498 | *pauca* | 20.17 (0.23) | 35.71 (1.51) |
| CFBP 8478 | *sandyi* | 19.81 (0.14) | na |
| CFBP 8356 | *sandyi* | 21.76 (0.17) | na |
| CFBP 8419 | *sandyi* | 22.04 (0.42) | na |

^a^: Mean Ct value performed on three independent tests. Results were considered as positive when at least six out of eight repetitions amplified the target DNA

^b^: not amplified

### Supplemental data 5: LOD of *Xf* in spiked matrices using the tetraplex qPCR assay XF – XFF – XFM – XFP (set n°2).

|  | **XF** | | **XFF** | | **XFM** | | **XFP** | | **Harper et al., 2010** | |
| --- | --- | --- | --- | --- | --- | --- | --- | --- | --- | --- |
| **Plant species**  **Spiked strain (subsp.)** | **LOD**  **(CFU.mL^-1^)** | **Mean Ct**  **(SEM)** | **LOD**  **(CFU.mL^-1^)** | **Mean Ct**  **(SEM)** | **LOD**  **(CFU.mL^-1^)** | **Mean Ct**  **(SEM)** | **LOD**  **(CFU.mL^-1^)** | **Mean Ct**  **(SEM)** | **LOD**  **(CFU.mL^-1^)** | **Mean Ct**  **(SEM)** |
| *Cistus monspeliensis* |  |  |  |  |  |  |  |  |  |  |
| CFBP 7970 (*fastidiosa*) | 1x10^5^ | 26.89 (0.45) | 1x10^4^ | 28.61 (1.45) |  | na^a^ |  | na | 1x10^2^ | 36.37 (0.06) |
| CFBP 8416 (*multiplex*) | 1x10^5^ | 27.46 (0.18) |  | na | 1x10^4^ | 28.11 (0.33) |  | na | 1x10^3^ | 36.48 (1.17) |
| *Citrus clementina* |  |  |  |  |  |  |  |  |  |  |
| CFBP 8402 (*pauca*) | 1x10^4^ | 27.47 (0.54) |  | na |  | na | 1x10^3^ | 29.05 (1.10) | 1x10^3^ | 31.72 (0.18) |
| CFBP 8416 (*multiplex*) | 1x10^3^ | 28.05 (0.21) |  | na | 1x10^3^ | 28.46 (0.10) |  | na | 1x10^2^ | 37.26 (0.20) |
| *Helichrysum italicum* |  |  |  |  |  |  |  |  |  |  |
| CFBP 8416 (*multiplex*) | 1x10^4^ | 27.52 (0.02) |  | na | 1x10^3^ | 31.00 (1.99) |  | na | 1x10^3^ | 32.96 (0.27) |
| *Lavandula angustifolia* |  |  |  |  |  |  |  |  |  |  |
| CFBP 8402 (*pauca*) | 1x10^4^ | 32.98 (2.87) |  | na |  | na | 1x10^4^ | 28.05 (0.07) | 1x10^3^ | 33.04 (0.20) |
| CFBP 8416 (*multiplex*) | 1x10^4^ | 30.37 (0.48) |  | na | 1x10^4^ | 28.55 (0.79) |  | na | 1x10^3^ | 33.71 (0.43) |
| *Nerium oleander* |  |  |  |  |  |  |  |  |  |  |
| CFBP 8402 (*pauca*) | 1x10^5^ | 27.09 (0.10) |  | na |  | na | 1x10^4^ | 28.51 (0.61) | 1x10^3^ | 35.86 (0.32) |
| CFBP 8416 (*multiplex*) | 1x10^5^ | 25.83 (0.02) |  | na | 1x10^4^ | 28.52 (0.24) |  | na | 1x10^3^ | 35.15 (0.31) |
| CFBP 8402 (*pauca*)  + CFBP 8416 (*multiplex*)^b^ | 1x10^6^ | 27.06 (0.12) |  | na | 5x10^5^ | 26.77 (1.03) | 5x10^5^ | 26.13 (0.18) | 1x10^3^ | 36.02 (0.31) |
| *Olea europaea* |  |  |  |  |  |  |  |  |  |  |
| CFBP 8402 (*pauca*) | 1x10^5^ | 25.54 (0.23) |  | na |  | na | 1x10^4^ | 26.35 (0.08) | 1x10^3^ | 33.71 (0.78) |
| CFBP 8416 (*multiplex*) | 1x10^5^ | 25.17 (0.09) |  | na | 1x10^5^ | 25.71 (0.14) |  | na | 1x10^3^ | 36.10 (1.35) |
| CFBP 8402 (*pauca*)  + CFBP 8416 (*multiplex*) | 1x10^6^ | 25.44 (0.17) |  | na | 5x10^5^ | 26.27 (0.10) | 5x10^5^ | 25.40 (0.19) | 1x10^6^ | 32.26 (0.17) |
| *Polygala myrtifolia* |  |  |  |  |  |  |  |  |  |  |
| CFBP 7970 (*fastidiosa*) | 1x10^5^ | 29.07 (1.22) | 1x10^5^ | 27.77 (0.20) |  | na |  | na | 1x10^3^ | 37.47 (0.29) |
| CFBP 8402 (*pauca*) | 1x10^5^ | 25.86 (0.05) |  | na |  | na | 1x10^4^ | 27.71 (0.36) | 1x10^3^ | 33.85 (0.43) |
| CFBP 8416 (*multiplex*) | 1x10^5^ | 26.75 (0.45) |  | na | 1x10^4^ | 30.05 (0.51) |  | na | 1x10^3^ | 33.17 (0.18) |
| CFBP 7970 (*fastidiosa*)  + CFBP 8402 (*pauca*)  + CFBP 8416 (*multiplex*) | 1x10^5^ | 26.08 (0.22) | 3.33x10^5^ | 23.14 (5.41) | 3.33x10^4^ | 27.49 (0.20) | 3.33x10^4^ | 25.14 (0.17) | 1x10^3^ | 36.23 (0.17) |
| *Prunus cerasus* |  |  |  |  |  |  |  |  |  |  |
| CFBP 7970 (*fastidiosa*) | 1x10^6^ | 26.78 (0.09) | 1x10^5^ | 26.85 (1.85) |  | na |  | na | 1x10^3^ | 35.69 (0.02) |
| CFBP 8416 (*multiplex*) | 1x10^6^ | 29.15 (0.15) |  | na | 1x10^6^ | 28.96 (0.11) |  | na | 1x10^4^ | 36.42 (0.45) |
| *Prunus dulcis* |  |  |  |  |  |  |  |  |  |  |
| CFBP 7970 (*fastidiosa*) | 1x10^5^ | 29.10 (0.29) | 1x10^5^ | 29.58 (1.25) |  | na |  | na | 1x10^4^ | 34.65 (0.96) |
| CFBP 8402 (*pauca*) | 1x10^5^ | 28.30 (0.25) |  | na |  | na | 1x10^4^ | 30.58 (0.05) | 1x10^4^ | 35.08 (0.15) |
| CFBP 8416 (*multiplex*) | 1x10^6^ | 27.83 (0.07) |  | na | 1x10^5^ | 30.07 (0.28) |  | na | 1x10^4^ | 37.17 (1.05) |
| CFBP 7970 (*fastidiosa*)  + CFBP 8402 (*pauca*)  + CFBP 8416 (*multiplex*) | 1x10^6^ | 33.36 (2.04) |  | na |  | na | 3.33x10^5^ | 27.06 (0.07) | 1x10^4^ | 35.89 (0.05) |
| *Quercus ilex* |  |  |  |  |  |  |  |  |  |  |
| CFBP 8416 (*multiplex*) | 1x10^5^ | 23.94 (0.06) |  | na | 1x10^4^ | 28.70 (1.02) |  | na | 1x10^2^ | 36.26 (1.34) |
| *Quercus robur* |  |  |  |  |  |  |  |  |  |  |
| CFBP 8416 (*multiplex*) | 1x10^5^ | 26.77 (0.08) |  | na | 1x10^5^ | 27.11 (0.03) |  | na | 1x10^3^ | 37.05 (0.09) |
| *Rosmarinus officinalis* |  |  |  |  |  |  |  |  |  |  |
| CFBP 8416 (*multiplex*) | 1x10^4^ | 27.84 (0.43) |  | na | 1x10^4^ | 27.58 (0.32) |  | na | 1x10^3^ | 32.55 (0.28) |
| *Vitis vinifera* |  |  |  |  |  |  |  |  |  |  |
| CFBP 7970 (*fastidiosa*) | 1x10^3^ | 30.27 (0.23) | 1x10^3^ | 26.44 (0.36) |  | na |  | na | 1x10^2^ | 37.65 (0.00) |
| CFBP 8416 (*multiplex*) | 1x10^3^ | 27.46 (2.03) |  | na | 1x10^3^ | 27.14 (1.62) |  | na | 1x10^3^ | 33.41 (0.19) |

^a^: not amplified

^b^: + indicates a mix infection by at least two strains

### Supplemental data 6: LOD of *Xf* in spiked matrices using the tetraplex qPCR assay XF – XFF – XFM – XFMO (set n°3).

|  | **XF** | | **XFF** | | **XFM** | | **XFMO** | | **Harper et al., 2010** | |
| --- | --- | --- | --- | --- | --- | --- | --- | --- | --- | --- |
| **Plant species**  **Spiked strain (subsp.)** | **LOD**  **(CFU.mL^-1^)** | **Mean Ct**  **(SEM)** | **LOD**  **(CFU.mL^-1^)** | **Mean Ct**  **(SEM)** | **LOD**  **(CFU.mL^-1^)** | **Mean Ct**  **(SEM)** | **LOD**  **(CFU.mL^-1^)** | **Mean Ct**  **(SEM)** | **LOD**  **(CFU.mL^-1^)** | **Mean Ct**  **(SEM)** |
| *Cistus monspeliensis* |  |  |  |  |  |  |  |  |  |  |
| CFBP 7970 (*fastidiosa*) | 1x10^5^ | 26.30 (0.15) | 1x10^4^ | 28.22 (1.39) |  | na^a^ |  | na | 1x10^2^ | 36.37 (0.06) |
| CFBP 8416 (*multiplex*) | 1x10^5^ | 28.35 (0.92) |  | na | 1x10^5^ | 27.61 (0.07) |  | na | 1x10^3^ | 36.48 (1.17) |
| *Citrus clementina* |  |  |  |  |  |  |  |  |  |  |
| CFBP 8416 (*multiplex*) | 1x10^4^ | 26.74 (0.01) |  | na | 1x10^3^ | 29.13 (0.43) |  | na | 1x10^2^ | 37.26 (0.20) |
| *Helichrysum italicum* |  |  |  |  |  |  |  |  |  |  |
| CFBP 8416 (*multiplex*) | 1x10^4^ | 27.64 (0.10) |  | na | 1x10^3^ | 28.64 (1.16) |  | na | 1x10^3^ | 32.96 (0.27) |
| *Lavandula angustifolia* |  |  |  |  |  |  |  |  |  |  |
| CFBP 8416 (*multiplex*) | 1x10^4^ | 27.67 (0.91) |  | na | 1x10^4^ | 28.87 (0.71) |  | na | 1x10^3^ | 33.71 (0.43) |
| *Nerium oleander* |  |  |  |  |  |  |  |  |  |  |
| CFBP 8416 (*multiplex*) | 1x10^5^ | 25.89 (0.02) |  | na | 1x10^4^ | 28.17 (0.12) |  | na | 1x10^3^ | 35.15 (0.31) |
| CFBP 8402 (*pauca*)  + CFBP 8416 (*multiplex*)^b^ | 1x10^4^ | 28.85 (0.09) |  | na | 5x10^4^ | 27.30 (0.14) |  | na | 1x10^3^ | 36.02 (0.31) |
| *Olea europaea* |  |  |  |  |  |  |  |  |  |  |
| CFBP 8416 (*multiplex*) | 1x10^5^ | 25.12 (0.08) |  | na | 1x10^5^ | 25.50 (0.15) |  | na | 1x10^3^ | 36.10 (1.35) |
| CFBP 8402 (*pauca*)  + CFBP 8416 (*multiplex*) | 1x10^6^ | 26.48 (0.06) |  | na | 5x10^5^ | 26.54 (0.13) |  | na | 1x10^6^ | 32.26 (0.17) |
| *Polygala myrtifolia* |  |  |  |  |  |  |  |  |  |  |
| CFBP 7970 (*fastidiosa*) | 1x10^5^ | 27.53 (0.66) | 1x10^5^ | 29.89 (1.43) |  | na |  | na | 1x10^3^ | 37.47 (0.29) |
| CFBP 8084 (*morus*) | 1x10^4^ | 27.91 (0.10) |  | na |  | na | 1x10^3^ | 26.09 (0.41) | 1x10^3^ | 32.53 (0.44) |
| CFBP 8416 (*multiplex*) | 1x10^4^ | 27.82 (0.15) |  | na | 1x10^3^ | 29.71 (1.93) |  | na | 1x10^3^ | 33.17 (0.18) |
| CFBP 7970 (*fastidiosa*)  + CFBP 8402 (*pauca*)  + CFBP 8416 (*multiplex*) | 1x10^5^ | 26.20 (0.12) | 3.33x10^5^ | 27.26 (0.26) | 3.33x10^4^ | 27.40 (0.16) |  | na | 1x10^3^ | 36.23 (0.17) |
| *Prunus cerasus* |  |  |  |  |  |  |  |  |  |  |
| CFBP 7970 (*fastidiosa*) | 1x10^6^ | 26.71 (0.05) | 1x10^5^ | 27.61 (0.51) |  | na |  | na | 1x10^3^ | 35.69 (0.02) |
| CFBP 8416 (*multiplex*) | 1x10^6^ | 28.73 (0.10) |  | na | 1x10^5^ | 30.59 (0.71) |  | na | 1x10^4^ | 36.42 (0.45) |
| *Prunus dulcis* |  |  |  |  |  |  |  |  |  |  |
| CFBP 7970 (*fastidiosa*) | 1x10^5^ | 29.17 (0.38) | 1x10^4^ | 31.30 (2.71) |  | na |  | na | 1x10^4^ | 34.65 (0.96) |
| CFBP 8416 (*multiplex*) | 1x10^6^ | 27.89 (0.39) |  | na | 1x10^5^ | 30.16 (0.52) |  | na | 1x10^4^ | 37.17 (1.05) |
| CFBP 7970 (*fastidiosa*)  + CFBP 8402 (*pauca*)  + CFBP 8416 (*multiplex*) | 1x10^5^ | 28.83 (0.16) | 1x10^5^ | 29.07 (0.22) | 1x10^6^ | 28.77 (0.04) |  | na | 1x10^4^ | 35.89 (0.05) |
| *Quercus ilex* |  |  |  |  |  |  |  |  |  |  |
| CFBP 8416 (*multiplex*) | 1x10^5^ | 23.85 (0.04) |  | na | 1x10^4^ | 27.00 (0.44) |  | na | 1x10^2^ | 36.26 (1.34) |
| *Quercus robur* |  |  |  |  |  |  |  |  |  |  |
| CFBP 8416 (*multiplex*) | 1x10^5^ | 27.03 (0.12) |  | na | 1x10^4^ | 28.14 (0.48) |  | na | 1x10^3^ | 37.05 (0.09) |
| *Rosmarinus officinalis* |  |  |  |  |  |  |  |  |  |  |
| CFBP 8416 (*multiplex*) | 1x10^5^ | 24.68 (0.04) |  | na | 1x10^4^ | 27.32 (0.23) |  | na | 1x10^3^ | 32.55 (0.28) |
| *Vitis vinifera* |  |  |  |  |  |  |  |  |  |  |
| CFBP 7970 (*fastidiosa*) | 1x10^4^ | 27.26 (0.12) | 1x10^3^ | 28.77 (0.86) |  | na |  | na | 1x10^2^ | 37.65 (0.00) |
| CFBP 8416 (*multiplex*) | 1x10^4^ | 28.04 (0.13) |  | na | 1x10^3^ | 33.29 (0.84) |  | na | 1x10^3^ | 33.41 (0.19) |

^a^: not amplified

^b^: + indicates a mix infection by at least two strains

### Supplemental data 7: *Xf* detection in dilution ranges of spiked matrices using the tetraplex qPCR assay XF – XFFSL – XFM – XFP (set n°1)

Xslx file

### Supplemental data 8: Comparison of LOD of *Xf* subsp. *fastidiosa* strain CFBP 7970 using the multiplex sets n°1, n°4, n°5 and n°6.

|  |  | **Set n°1 XF-XFFSL-XFM-XFP** | |  | **Set n°4 XFFSL-XFM-XFP** |  | **Set n°5 Harper-XFFSL-XFM-XFP** | |  | **Set n°6 18S-XFFSL-XFM-XFP** | |
| --- | --- | --- | --- | --- | --- | --- | --- | --- | --- | --- | --- |
| **Concentration** | **Theoretical**  **number of**  **copy.mL^-1^** | **XF mean Ct (SEM)** | **XFFSL mean Ct (SEM)** |  | **XFFSL mean Ct (SEM)** |  | **Harper mean Ct (SEM)** | **XFFSL Mean Ct (SEM)** |  | **18S mean Ct (SEM)** | **XFFSL mean Ct (SEM)** |
| 1 µg.mL^-1^ | 4x10^8^ | 17.59 (0.03) | 18.49 (0.06) |  | 18.53 (0.08) |  | 18.12 (0.01) | 18.18 (0.02) |  | na | 18.51 (0.02) |
| 100 ng.mL^-1^ | 4x10^7^ | 20.98 (0.06) | 21.93 (0.02) |  | 22.04 (0.02) |  | 21.65 (0.04) | 21.75 (0.05) |  | na | 21.91 (0.02) |
| 10 ng.mL^-1^ | 4x10^6^ | 23.92 (0.04) | 24.81 (0.02) |  | 25.82 (0.02) |  | 25.41 (0.12) | 25.56 (0.14) |  | na | 25.81 (0.07) |
| 1 ng.mL^-1^ | 4x10^5^ | 26.48 (0.10) | 27.51 (0.12) |  | 28.27 (0.03) |  | 27.76 (0.08) | 28.01 (0.02) |  | na | 28.10 (0.04) |
| 100 pg.mL^-1^ | 4x10^4^ | 27.79 (0.33) | 30.24 (0.11) |  | 30.90 (0.17) |  | 30.47 (0.42) | 31.03 (0.57) |  | na | 30.30 (0.26) |
| 10 pg.mL^-1^ | 4x10^3^ | na^a^ | 31.81 (0.13) |  | 31.25 (0.98) |  | 35.93 (1.94) | 31.08 (0.53) |  | na | 32.06 (1.37) |
| 1 pg.mL^-1^ | 4x10^2^ | na | na |  | na |  | na | na |  | na | na |

^a^: not amplified

### Supplemental data 9: Comparison of LOD of *Xf* in spiked matrices using the multiplex set n°1, n°4, n°5 and n°6.

|  |  | **Set n°1 XF-XFFSL-XFM-XFP** | | **Set n°4 XFFSL-XFM-XFP** | **Set n°5 Harper-XFFSL-XFM-XFP** | | **Set n°6 18S-XFFSL-XFM-XFP** | |
| --- | --- | --- | --- | --- | --- | --- | --- | --- |
| **Plant species**  **Spiked strain (Xf subspecies)** | **Theoretical**  **number of**  **CFU.mL^-1^** | **XF Mean Ct (SEM)** | **XFM Mean Ct (SEM)** | **XFM Mean Ct (SEM)** | **Harper Mean Ct (SEM)** | **XFM Mean Ct (SEM)** | **18S Mean Ct (SEM)** | **XFM Mean Ct (SEM)** |
| *Q. robur*  CFBP 8416 (*multiplex*) |  |  |  |  |  |  |  |  |
|  | 1x10^6^ | 24.41 (0.05) | 23.78 (0.03) | 23.92 (0.09) | 24.56 (0.23) | 23.89 (0.28) | 17.12 (0.03) | 24.06 (0.04) |
|  | 1x10^5^ | 26.85 (0.04) | **26.16 (0.14)** | **26.40 (0.23)** | 27.15 (0.10) | **26.48 (0.11)** | 17.54 (0.08) | **26.80 (0.01)** |
|  | 1x10^4^ | **28.99 (0.27)^a^** | na | na | **30.92 (0.48)** | na | 18.14 (0.03) | na |
|  | 1x10^3^ | na^b^ | na | na | na | na | 17.89 (0.11) | na |
|  | 1x10^2^ | na | na | na | na | na | 18.22 (0.01) | na |
|  | 1x10^1^ | na | na | na | na | na | 18.56 (0.06) | na |
|  | T- | na | na | na | na | na | **18.33 (0.08)** | na |
| *C. monspeliensis*  CFBP 8416 (*multiplex*) |  |  |  |  |  |  |  |  |
|  | 1x10^6^ | 26.61 (0.02) | 25.87 (0.01) | 26.62 (0.06) | 26.53 (0.14) | 25.87 (0.09) | 20.97 (0.05) | 26.46 (0.10) |
|  | 1x10^5^ | **27.14 (0.09)** | **26.55 (0.13)** | **26.94 (0.09)** | 27.48 (0.06) | **26.89 (0.03)** | 19.50 (0.07) | **26.97 (0.06)** |
|  | 1x10^4^ | na | na | na | **32.10 (1.39)** | na | 20.33 (0.07) | na |
|  | 1x10^3^ | na | na | na | na | na | 19.71 (0.05) | na |
|  | 1x10^2^ | na | na | na | na | na | 19.77 (0.08) | na |
|  | 1x10^1^ | na | na | na | na | na | 20.26 (0.24) | na |
|  | T- | na | na | na | na | na | **19.05 (0.06)** | na |

^a^: bold text: threshold retained when at least two out of three replicates amplified

^b^: not amplified
